# Functional architecture of the synaptic transducers at a central glutamatergic synapse

**DOI:** 10.1101/2020.12.25.424391

**Authors:** Marisa M. Brockmann, Estelle Toulme, Andreas T. Grasskamp, Thorsten Trimbuch, Thomas C. Südhof, Alexander M. Walter, Christian Rosenmund

## Abstract

Neuronal synapses transduce information *via* the consecutive action of three transducers: voltage-gated Ca^2+^-channels, fusion-competent synaptic vesicles, and postsynaptic receptors. Their physical distance is thought to influence the speed and efficiency of neurotransmission. However, technical limitations have hampered resolving their nanoscale arrangement. Here, we developed a new method for live-labeling proteins for electron microscopy (EM), revealing that release-competent vesicles preferentially align with Ca^2+^-channels and postsynaptic AMPA receptors within 20-30 nm and thereby forming a transsynaptic tripartite nanocomplex. Using functional EM, we show that single action potentials cause vesicles within the nanocomplex to fuse with a 50% probability. The loss of the presynaptic scaffold disrupts the formation of the tripartite transducers. Strikingly, the forced transsynaptic alignment of the Ca^2+^-channel subunit α2δ1 and AMPA receptors suffice to restore neurotransmission in a scaffold lacking synapse. Our results demonstrate a synaptic transducer nanocomplex that actively contributes to the organization of central synapses.

## Introduction

Synapses are highly specialized structures that orchestrate the activity-dependent information transfer between neurons. At the presynapse, an incoming action potential opens voltage-gated Ca^2+^-channels (VGCC) that triggers the fusion of release competent synaptic vesicles (SVs) and neurotransmitter release into the synaptic cleft (Sudhof, 2014). At the postsynapse, the neurotransmitter is sensed by its respective receptors and elicits an amplitude-coded response. Therefore, Ca^2+^-channels, fusion-competent SVs, and postsynaptic receptors represent the key transducers of synaptic transmission. While this basic concept is well-accepted, a clearer picture of the nanoscale arrangement of pre- and postsynaptic proteins is only now starting to emerge.

The nanoscale arrangement of Ca^2+^-channels and fusion-competent SVs is thought to be critical in determining the speed and efficiency of neurotransmission. Initial experiments, performed at the frog neuromuscular junction (Harlow et al., 2001) and the squid giant synapse (Adler et al., 1991), already suggested tight coupling distances between Ca^2+^-channels and fusion-competent SVs that favor efficient Ca^2+^ secretion coupling. Measuring the sensitivity of neurotransmitter release in the presence of exogenous calcium chelators indicates a wide range (10 nm and 100 nm) of functional coupling distances at mammalian central synapses (Arai and Jonas, 2014; Bohme et al., 2016; Bucurenciu et al., 2008; Eggermann et al., 2011; Keller et al., 2015; Nakamura et al., 2015; Rebola et al., 2019; Vyleta and Jonas, 2014). However, the physical distance between fusion-competent SVs and Ca^2+^-channels in small mammalian central synapses has not yet been measured.

Distinct coupling distances between Ca^2+^-channels and release-competent SVs may contribute to the striking diversity of release probability and short-term plasticity characteristics among synapses (Nusser, 2018; Rosenmund et al., 1993). RIM proteins (Rab3-interacting molecule) and RIM-binding proteins (RBPs) are particularly good candidates to control the Ca^2+^-secretion coupling distance in synapses. They directly interact with Ca^2+^-channels, and their loss of function leads to disruption of Ca^2+^-secretion coupling in both vertebrates and invertebrates (Acuna et al., 2015; Brockmann et al., 2019; Davydova et al., 2014; Graf et al., 2012; Grauel et al., 2016; Kaeser et al., 2011; Liu et al., 2011). Therefore, by controlling the precise distances between fusion-competent SVs and voltage-gated Ca^2+^-channels, these presynaptic scaffolding proteins could contribute to the heterogeneity of release properties between synapses.

The positional arrangement of central postsynaptic glutamate receptors also shapes synaptic function. AMPA receptors are organized in nanodomains (MacGillavry et al., 2013), and super-resolution microscopy revealed a transsynaptic organization of nanocolumns, including the association of the presynaptic scaffold RIM with the postsynaptic scaffold PSD-95 and AMPA receptors (Tang et al., 2016). These findings raise the hypothesis that synapses comprise functional units at the pre- and postsynapse that must be transsynaptically aligned to ensure successful neurotransmission. Therefore, it is not only important to determine the spatial arrangement of functional units, but also to uncover the underlying molecular mechanism that drives their assembly. Although multiple transsynaptic interactions by diverse adhesion molecules contribute to the assembly of pre- and postsynaptic assemblies (Sudhof, 2018), we still lack information about which and how those interactions control the position of the key transducers.

In this study, we aimed to address the relative localization of the three key transducers of efficient neurotransmission — fusion-competent SVs, Ca^2+^-channels, and AMPA receptors — in the synapse. To determine the transducers’ spatial arrangement on a nanometer scale, we developed a live-gold-labeling, EM compatible tagging approach for synaptic proteins to reveal the spatial arrangement of docked SVs in the active zone relative to proteins, such as Ca^2+^-channels. We could detect Ca^2+^- and AMPA channels within the narrow confines of the synaptic cleft and found that they both align with a distance of 20-30 nm to docked SVs to form a nanocomplex in small excitatory synapses. Moreover, we combined the live-labeling of Ca^2+^-channels with functional EM, allowing us to arrest the synaptic ultrastructure milliseconds after action-potential induced SV fusion and defined the vesicle fusion probability as a function of Ca^2+^-channel distance. Synaptic vesicles within the nanocomplex fused with a 50% probability upon action potential stimulation. Furthermore, genetically induced the loss of synaptic scaffold led to loss of the alignment of the three transducers and synaptic transmission, but by forcing the transsynaptic alignment of the presynaptic voltage-gated Ca^2+^-channel auxiliary subunit α2δ1 and postsynaptic AMPAR receptor subunit GluA2, synaptic transmission was restored. Thus, α2δ1 can act as a nucleation factor to recruit the Ca^2+^-channels α-subunit and docked SVs to enable neurotransmission in the absence of a presynaptic scaffold. Therefore, our findings support a model of self-organizing synaptic units and close physical Ca^2+^-channel-fusion-competent-SV coupling as means by which central mammalian synapses ensure efficient synaptic transmission.

## Results

We first sought to localize voltage-gated Ca^2+^-channels within the presynaptic active zone (AZ). We tagged the mouse Ca^2+^-channel auxiliary subunit α2δ1 (Dolphin, 2013) with a 10 x His-Tag and induced its exogenous expression using lentivirus (Figure 1A). Using nickel nitrilotriacetic acid (NiNTA) coupled to 5 nm gold particles, we were able to detect α2δ1-His with a coupling distance of 1.5 nm within the narrow confines of the synaptic cleft.

**Figure 1:**
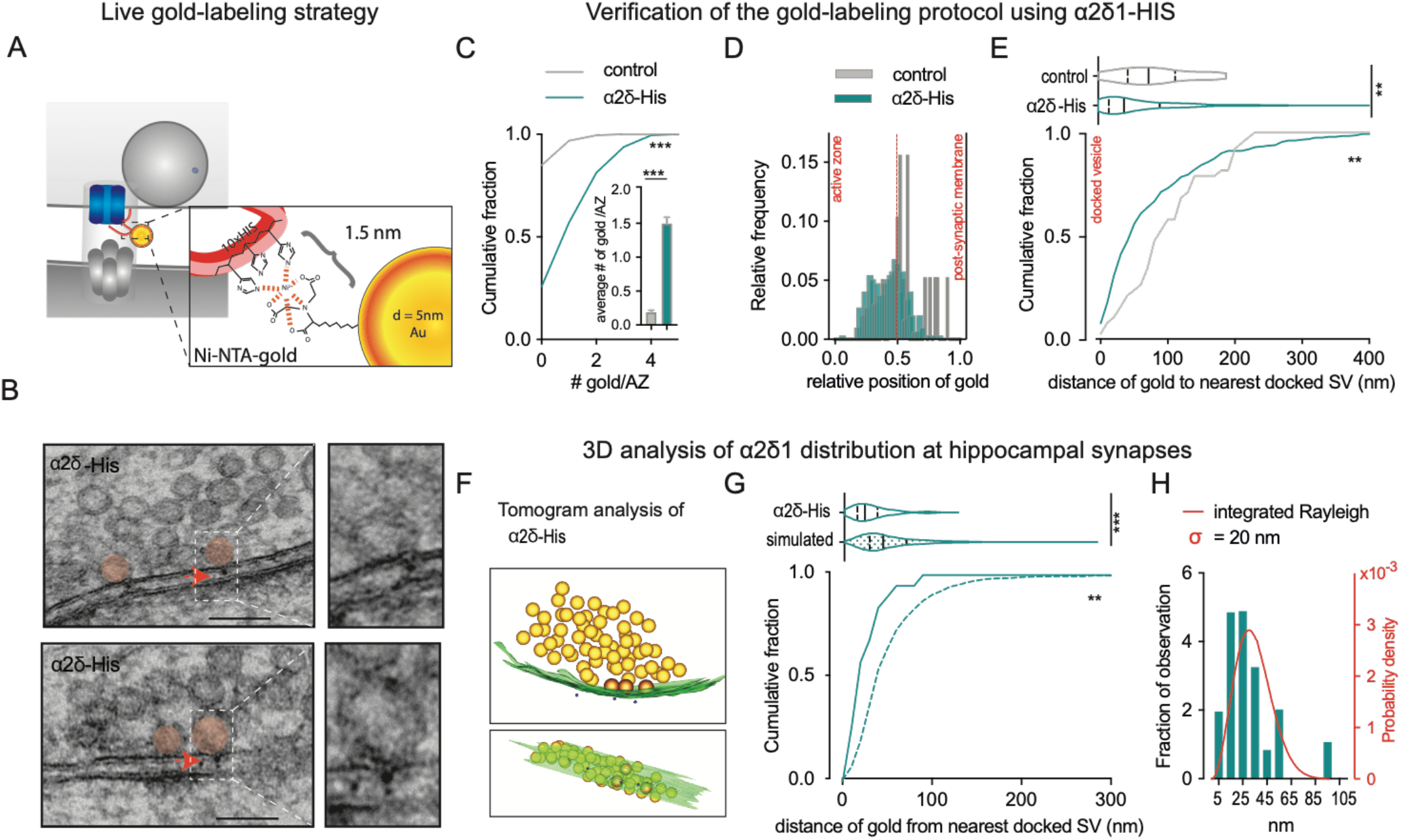
A novel tool to gold-label synaptic proteins reveal the tight association of docked SVs with Ca^2+^-channels. **(A**) Live gold-labeling strategy of the Ca^2+^-channel subunit α2δ using polyhistidine-tagging and incubation with 5 nm NiNTA-gold particles; coupling 1.5 nm. (**B**) Representative electron micrographs of cryofixed primary hippocampal neurons expressing α2δ1-His. Scale bars, 100 nm (docked SV = red circle, arrows point to gold particles). (**C**) Number of gold particles per AZ. (**D**) Gold particles distribution throughout the synaptic cleft width (0 = active zone membrane; 1 = post-synaptic membrane). (**E**) Distance from each gold particle to the nearest docked SV in a synaptic profile from control and α2δ1-His transfected neurons. (**F)** Tomogram of a cryofixed excitatory hippocampal synapse transfected with α2δ1-His (docked SV = orange, gold particle = blue, active zone = green). (**G)** Distance of each gold particle to the nearest docked SV in tomograms in α2δ1-His expressing neurons, compared to random placement. (**H)** Histogram of the distance of each gold particle to the nearest docked SV. Integrated Rayleigh distribution (red line). Data for bar graphs are means ± SEM. Data for violin plots are medians (solid lines) and quartiles (dashed lines). Statistical significance for C was assessed by unpaired t-test, the violin plots for E and G by Mann-Whitney test, and the cumulative distribution plots in C, E, and G by Kolmogorov-Smirnov test. *p < 0.05, **p < 0.01, *** p < 0.001.

Because of their synaptogenic function, strong overexpression of α2δ1 in neurons potentiates synaptic responses (Eroglu et al., 2009; Hoppa et al., 2012). To prevent alterations in synaptic localization of α2δ1-His due to its overexpression, we transduced α2δ1-His in primary neurons and assessed synaptic responses. The concentrations of α2δ1-His reached in our experiments did not affect synaptic amplitudes or release probability (Figure S1). Moreover, ultrastructural parameters of the synapse, such as the number and distribution of docked SV at the AZ and the length of the postsynaptic density (PSD), were also unchanged upon α2δ1-His expression (Figure S2A-E). Pre-incubation of synapses expressing α2δ1-His with the NiNTA-gold resulted in an 8-fold enrichment of gold particles in the synaptic cleft (Figure 1B, C), and their position was significantly skewed towards the presynaptic membrane within the synaptic cleft width (Figure 1D), indicating specific labeling. The gold particles were equally allocated across the length of the synaptic profile, suggesting a uniform distribution of the Ca^2+^-channel subunit α2δ1, similar to the positions of docked SVs (Figure S2E, I).

### Ca^2+^-channels associate with docked SVs in 20 nm

While physiological recordings, super-resolution imaging, and computational studies inferred the coupling distance in various vertebrate and invertebrate synapses (Arai and Jonas, 2014; Bucurenciu et al., 2008; Eggermann et al., 2011; Keller et al., 2015; Nakamura et al., 2015; Schmidt et al., 2013; Vyleta and Jonas, 2014; Wadel et al., 2007), the distance of Ca^2+^-channels to fusion-competent, docked SVs has never been directly determined. We assessed each gold particle’s shortest distance, labeling α2δ1, to the next docked SV. The cumulative distribution, as well as the mean-nearest neighbor analysis, revealed that the distance between the Ca^2+^-channel subunit α2δ1 and the next docked SV is non-random and significantly different from the control, skewing the distribution towards shorter distances (Figure 1E, and Figure S2J). In the first 60 nm of the next docked SV, the abundance of α2δ1-His increased by 23 % compared to simulations of uniform SV and gold particle placement (Figure S2L), with a median distance of 46 nm (Figure 1E). To gain further insight into the Ca^2+^-channel-docked SV relationship at the AZ, we obtained tomograms and analyzed the distance of α2δ1-His to the center of the next docked SV in 3D (Figure 1F, G, Figure S3, and Figure S4A). Strikingly, as with the 2D analysis, the 3D analysis revealed an accumulation of α2δ1 within a median distance of 22 nm of the closest docked SV compared to random uniform SV and gold particle placement (Figure 1G, and Figure S4B). Fitting the minimal distances, using an integrated Rayleigh function, revealed a *σ* value of 20.3 nm (Figure 1H), demonstrating a close association of Ca^2+^-channels with docked SVs.

### The vesicle release probability as a function of Ca^2+^-channels to fusion-competent SV distance

In theory, the tight association of Ca^2+^-channel and fusion competent SVs should promote the high probability of action potential triggered SV fusion (Augustine et al., 2003; Bucurenciu et al., 2008; Neher, 1998). However, how this translates into absolute release probabilities as a function of the distance of two key presynaptic transducers and how release probabilities of docked SVs decay with increasing distances has only been modeled. We set out to define the probability of SVs fusion as a function of Ca^2^-channel distance by comparing the distances of Ca^2+^-channels and fusion-competent SVs between unstimulated samples and samples arrested milliseconds after action potential induced vesicle fusion (Figure 2A). This precise timing was now possible by combining our novel anatomical approach with stim-and-freeze technology, in which a single action potential is triggered in neurons by field stimulation, followed by rapid cryofixation with a delay of 5 ms using a high-pressure freezing device. The number of docked SVs in stimulated sections decreased by 30 %, demonstrating successful fusion events across the whole AZ length (Figure 2B and Figure S2C, E) (Kusick et al., 2020). As predicted, the median distance of α2δ1-His to the next remaining docked SV increased from 46 nm to 126 nm since docked SVs close to Ca^2+^-channels had been depleted. These findings strongly support our assumption that synaptic α2δ1 labeling is a proxy for functional Ca^2+^-channels (Figure 2C). Therefore, we were now able to define the vesicular release probability as a function of distances to Ca^2+^-channels. By comparing the minimal distances between gold particles and SVs without stimulation to the ones observed 5 ms after stimulation, we found a 50% vesicular release probability at a distance of 20 nm (Figure 2D). The likelihood of SV fusion as a function of distance was determined by fitting with a single exponential decay function and revealed a peak release probability of 86% for minimal distances while release probability decayed with a length constant of 39 nm. This value represents an upper limit of the Ca^2+^-secretion coupling distance as live labeling of Ca^2+^-channels by gold particles was likely not saturated.

**Figure 2:**
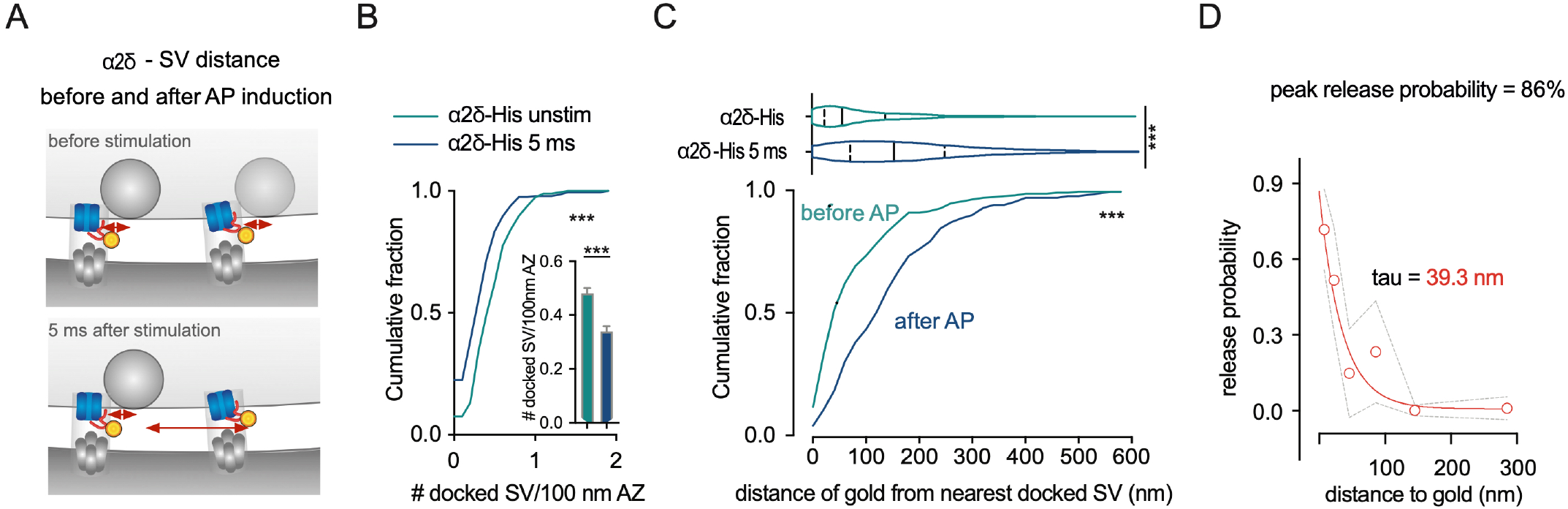
Deriving SV release probability from Ca^2+^ channel distance. **(A)** Experimental design to determine the distance of Ca^2+^-channels to docked SVs before and after the induction of an action potential. To obtain SVs’ distribution after action potential stimulation, the synaptic ultrastructure was arrested 5 ms after stimulation. (**B)** Docked SVs per synaptic profile in α2δ1-His transfected neurons cryo-fixed before (green lines, bars) and after (blue lines, bars) stimulation. (**C)** Distance of each gold particle to the nearest docked SV in a synaptic profile, comparing unstimulated and stimulated conditions. (**D)** Calculation of the vesicular release probability as a function of distances to α2δ1 gold label. Data for bar graphs are means ± SEM. Data for violin plots are medians (solid lines) and quartiles (dashed lines). Statistical significance for B was assessed by unpaired t-test, the violin plot for C by Mann-Whitney test, and the cumulative distribution plots in B and C by Kolmogorov-Smirnov test. ***p < 0.001.

### AMPA receptors transsynaptically align with docked SVs

The magnitude of the postsynaptic response to presynaptic neurotransmitter release is defined both by the number of postsynaptic receptors (Nusser et al., 1998) and the fraction of receptor binding sites occupied by glutamate; the latter of which increases with close apposition to the site of presynaptic release (Savtchenko and Rusakov, 2014). Therefore, we tested whether AMPA receptors are also associated with docked SVs by labeling exogenously expressed His-tagged AMPA receptor GluA2 subunits in living neurons (Figure 3A, B). As with α2δ1-His expression, GluA2-His expression did not affect the number and distribution of docked SVs at the AZ or the PSD length (Figure S5A-E). We detected a ∼5-fold enrichment of gold particles in the synaptic cleft upon GluA2-His expression, with localization skewed towards the postsynaptic membrane within the width of the synaptic cleft (Figure S5F-H). Interestingly, the synaptic distribution of GluA2 showed a preference towards the edge of the AZ (Figure 3C). To determine whether GluA2 is preferentially localized near release sites, we measured each gold particle’s distance to the nearest presynaptic docked SV. Again, we found a non-random distribution with a 57 % higher probability of GluA2-His within 60 nm of docked SVs compared to randomized data (Figure 3D, E). Strikingly, in the first 90 nm, the cumulative distribution of GluA2-His to docked SVs is equal to the distribution of α2δ1-His (Figure 3D). These data compellingly suggest that the three main transducers of synaptic transmission constitute an anatomical unit.

**Figure 3:**
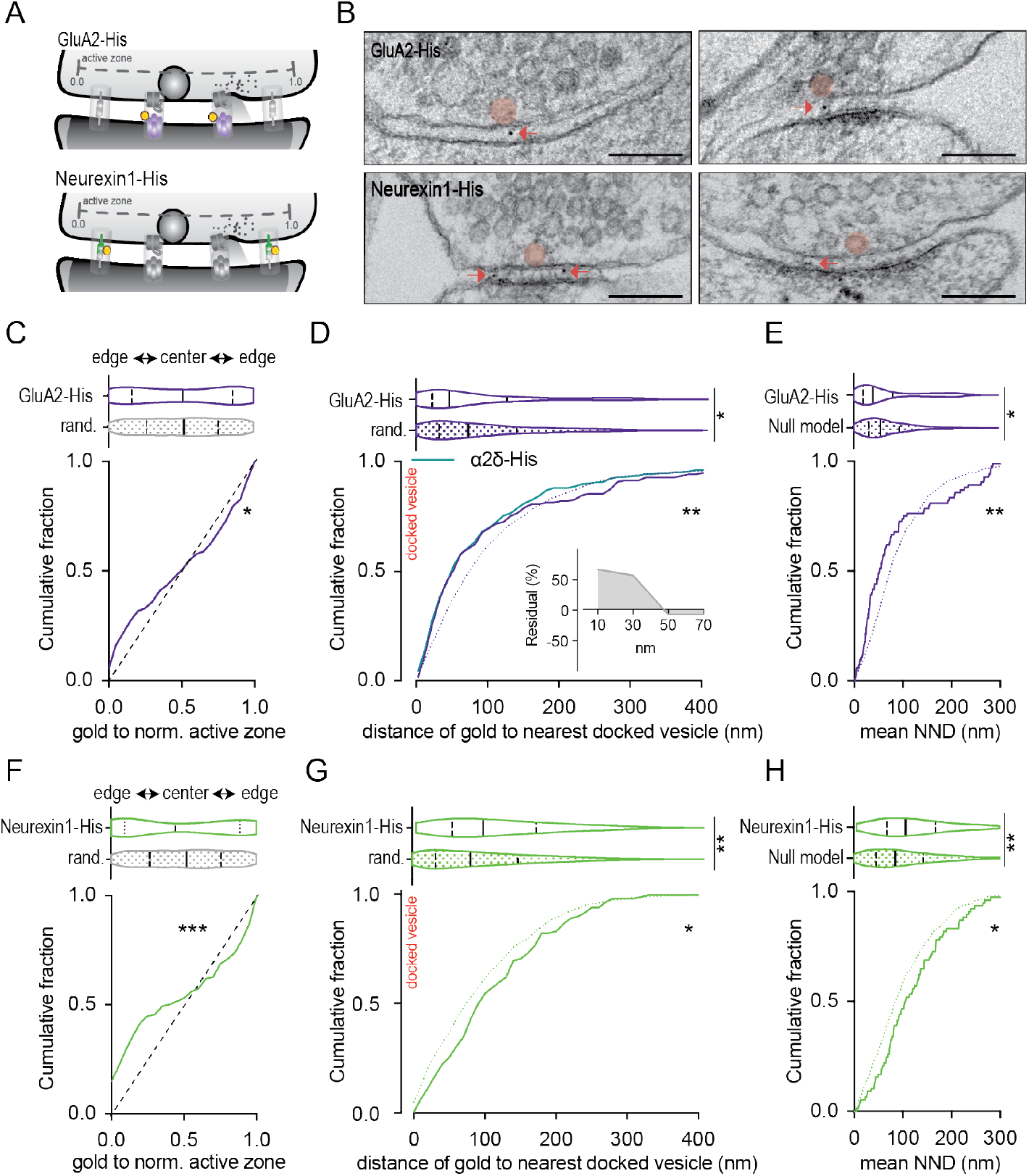
GluA2, but not neurexin1, is associated with docked SVs. **(A)** Graphic for the ultrastructural parameters analyzed in electron micrographs for GluA2-His and neurexin1-His expressing neurons. (**B)** Representative electron micrographs of cryofixed primary hippocampal neurons expressing GluA2-His or neurexin1-His. (**C and E)** Violin, and cumulative probability plots of gold particle distribution at the normalized AZ. (**D and G)** Violin, and cumulative probability plots of the nearest distance between gold participles and the next docked SV compared to randomized distances (dashed line). Insert depicts the residual percentage of the distance of gold particles to the next docked SV compared to their randomized distribution. (**E and H)** Violin, and cumulative probability plots of the measured data and null model data of mean nearest neighbor distance (NND) to docked SVs. Data for violin plots are medians (solid lines) and quartiles (dashed lines). Statistical significance for violin plots in C, D, E, F, G, and H was assessed by the Mann-Whitney test and for the cumulative distribution plots C, D, E, F, G, and H by Kolmogorov-Smirnov test. *p < 0.05, **p < 0.01, *** p < 0.001.

### Neurexin1 and neuroligin1 are localized at the edge of the AZ

At mammalian hippocampal synapses, pre- and postsynaptic scaffold proteins align in nanocolumns across the synapse, possibly mediated by cell-adhesion molecules (Biederer et al., 2017; Sudhof, 2017). Transsynaptic alignment of pre- and postsynaptic scaffold proteins has been linked to adhesion molecules (Sudhof, 2018). To test for their putative role in positioning Ca^2+^-channels, release competent SVs, and postsynaptic receptors, we expressed His-tagged neurexin1 and neuroligin1 in neurons and probed their synaptic distribution. Neurexin1-His and neuroligin1-His expression did not alter synaptic ultrastructure (Figure S5A-E) and resulted in an ∼5-fold enrichment of gold particles in the synaptic cleft. Gold particles were closer to the presynapse for neurexin1 and closer to the postsynapse for neuroligin1 (Figure S5F-H), consistent with their predicted pre- and postsynaptic localization, respectively (Sudhof, 2017). Along the surface of the AZ, neuroligin1, and to a greater extent, neurexin1 show a non-randomized distribution, with a strong enrichment towards the edge of the AZ (Figure 3F and Figure S5I). When assessing their relative position to docked SVs, neurexin1 and neuroligin1 showed an increased distance to docked SV compared to randomized placement (Figure 3G, H and Figure S5J, K). Therefore, neurexin1 and neuroligin1 are not closely positioned to docked SVs and are unlikely to drive the tripartite synaptic transducer complex assembly.

### The presynaptic scaffold orchestrates the transducers’ alignment

So far, our EM data show that Ca^2+^-channels, release competent SVs, and postsynaptic AMPA receptors form an anatomical unit. The main mechanisms proposed how the transducers align at the AZ is *via* synaptic scaffold proteins that align in nanocolumns and subsequently recruit and stabilize them. We addressed the synaptic scaffold’s requirement by examining synapses from the RIM/RBP quadruple KO (qKO) mouse. In qKO neurons, the presynaptic scaffold is drastically disrupted, docked SVs, and presynaptic Ca^2+^-channels are substantially reduced, the PSD length is increased, and consequently, synaptic transmission nearly abolished (Figure S6) (Acuna et al., 2016; Brockmann et al., 2020). Our live-gold-labeling protocol showed that in qKO synapses, His-tagged α2δ1 and GluA2 were significantly reduced (Figure 4A-D). Consistent with the electron microscopy data, dSTORM analysis revealed a severe, synapse-specific loss of endogenous Cav2.1, Cav2.2, and GluA1 and GluA2 protein clusters (Figure 4F-H, and Figure S6F-O). The reduction in postsynaptic AMPA receptors was confirmed by electrophysiological recordings detecting a 60 % reduction in the mEPSC amplitude (Figure 4I, J). Interestingly, mEPSC rise time in qKO neurons was significantly increased, perhaps caused by the broadening of the synaptic cleft width or the aberrant alignment of fusing SVs to postsynaptic receptors (Figure 4E, K). The mEPSC amplitude and rise time were obtained from the reanalysis of previously published data (Brockmann et al., 2020). These findings strongly support that the synaptic scaffold is necessary for the efficient alignment of the tripartite transducer complex. However, there is emerging evidence that the transducers transsynaptically align by direct or indirect interactions (Watson et al., 2017), and it is possible that the transducers autonomously promote their coupling and therefore promote synaptic transmission. To test this, we bypassed scaffold function by the forced alignment of two transducers, α2δ1 and GluA2, in a scaffold lacking synapse and tested for their independent role in synaptic organization and function.

**Figure 4:**
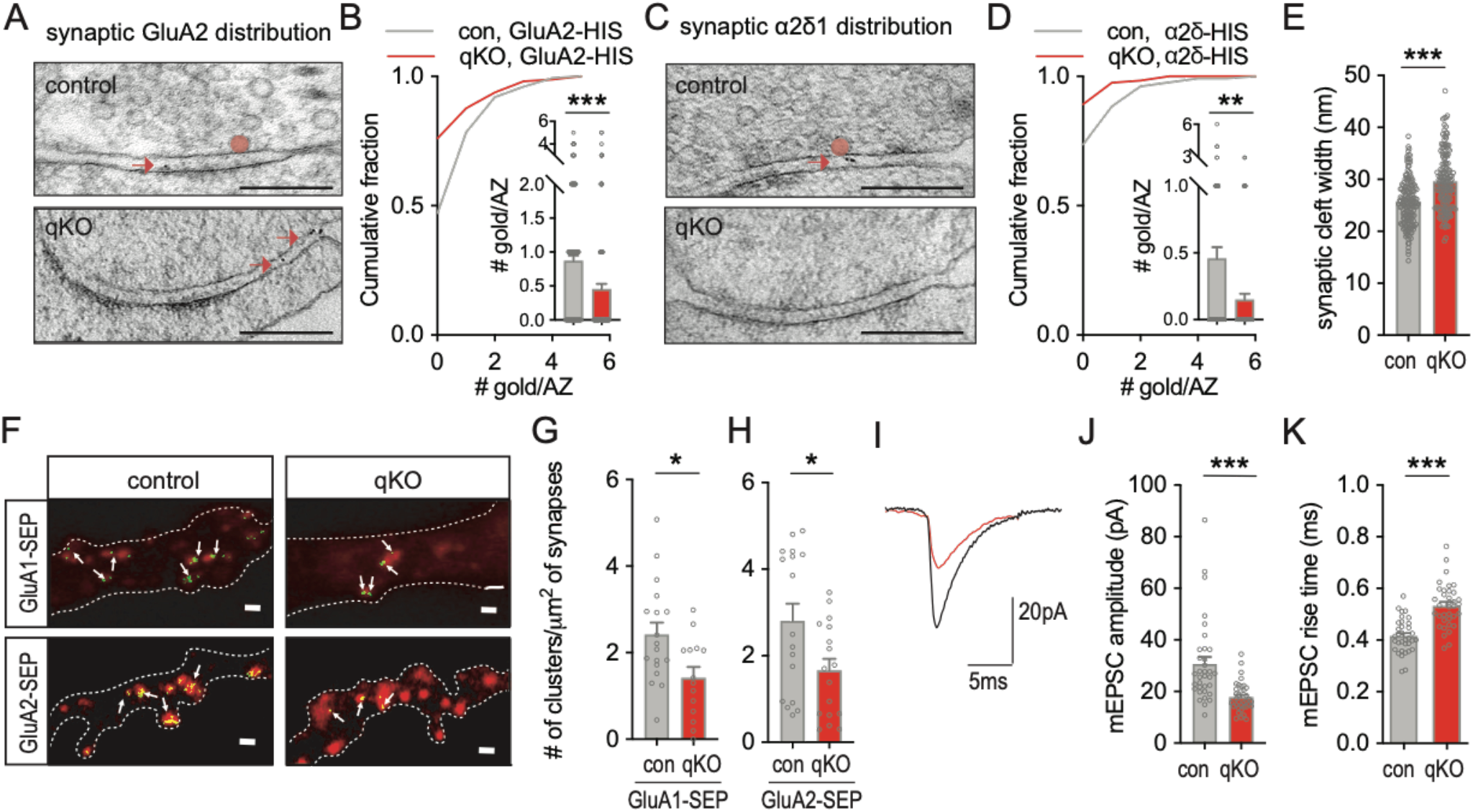
Loss of transducer-coupling in RIM/RBP deficient synapses. **(A-H)**, Experiments were conducted in RIM/RBP deficient synapses (qKO) and their respective control (Δcre). (**A and C)** Representative electron micrographs of cryo-fixed control and qKO primary hippocampal neurons expressing either GluA2-His or α2δ1-His. Scale bar, 200 nm. (**B and D)** Number of gold particles per AZ. **(E)** Synaptic cleft width. (**F)** Representative dSTORM images of GluA1 and GluA2 (green) and Homer (red) expressed in control and qKO neurons. Scale bars 1 µm. (**G)** Number of GluA1 clusters per synapse area. (**H)** Number of GluA2 clusters per synapse area. (**I)** Sample traces of mEPSC events in control (grey) and qKO (red) derived from autaptic neurons. (**J)** Miniature excitatory postsynaptic current (mEPSC) amplitude. (**K)** mEPSC rise time. Data are individual values and means ± SEM. Statistical significance for B, D, E, J, K, G, and H was assessed by unpaired t-test and cumulative distribution plots in B and D by Kolmogorov-Smirnov test. *p < 0.05, ***p < 0.001.

### Artificial alignment of the transducers’ rescues neurotransmission in a scaffold lacking synapse

We reconstituted Ca^2+^-channel and AMPA receptor alignment in qKO synapses by artificially enforcing the extracellular interaction of α2δ1- and GluA2 through a transsynaptic biotin-minimal streptavidin (mSA) link (Penn et al., 2017). We then tested whether this link would, in turn, restore some synaptic architecture and function. To our surprise, transfecting α2δ1-acceptor protein (AP), GluA2-mSA, and the biotin ligase BirA in qKO synapses (Figure 5A) rescued synaptic transmission (Figure 5B, C). Multiple functions are required for synaptic transmission. First, to test for the reassembly of release sites, we probed for the formation of plasma membrane docked, fusion-competent SVs using EM. We found that SV docking, and its electrophysiology equivalent, SV priming, were robustly rescued (Figure 5D-G), likely due to partial relocalization of the essential SV priming factor Munc13-1 that was lost in qKO synapses (Figure S7H, J) (Acuna et al., 2016; Brockmann et al., 2020). Second, voltage-gated Ca^2+^-channel function returned in the presynaptic terminals of qKOs neurons (Figure 5H, I), indicative for the recruitment of pore-forming Ca^2+^-channel α-subunits to release sites. The absence of neurotransmission in synapses, only expressing α2δ1-AP and GluA2-mSA without BirA, where no interaction between the two components was forced, shows that not the re-expression itself but specifically their alignment in the synapse is mandatory for neurotransmission. Overall, these data suggest that the stable alignment of Ca^2+^-channels, AMPA receptors, and docked SV is both necessary and sufficient to achieve neurotransmission. Additionally, anchoring α2δ1 at the presynapse is a key step in organizing the presynaptic release function. However, not all functions lost in qKO synapses were restored by the artificial alignment of α2δ1 and GluA2: the increase in PSD length and synaptic cleft width remained, and also mEPSC amplitude and rise time were not significantly rescued (Figure S7A-G). These data suggest that the transsynaptic Ca^2+^-channel-GluA2 alignment restored presynaptic release, it did not recruit endogenous AMPA channels across functional release sites.

**Figure 5:**
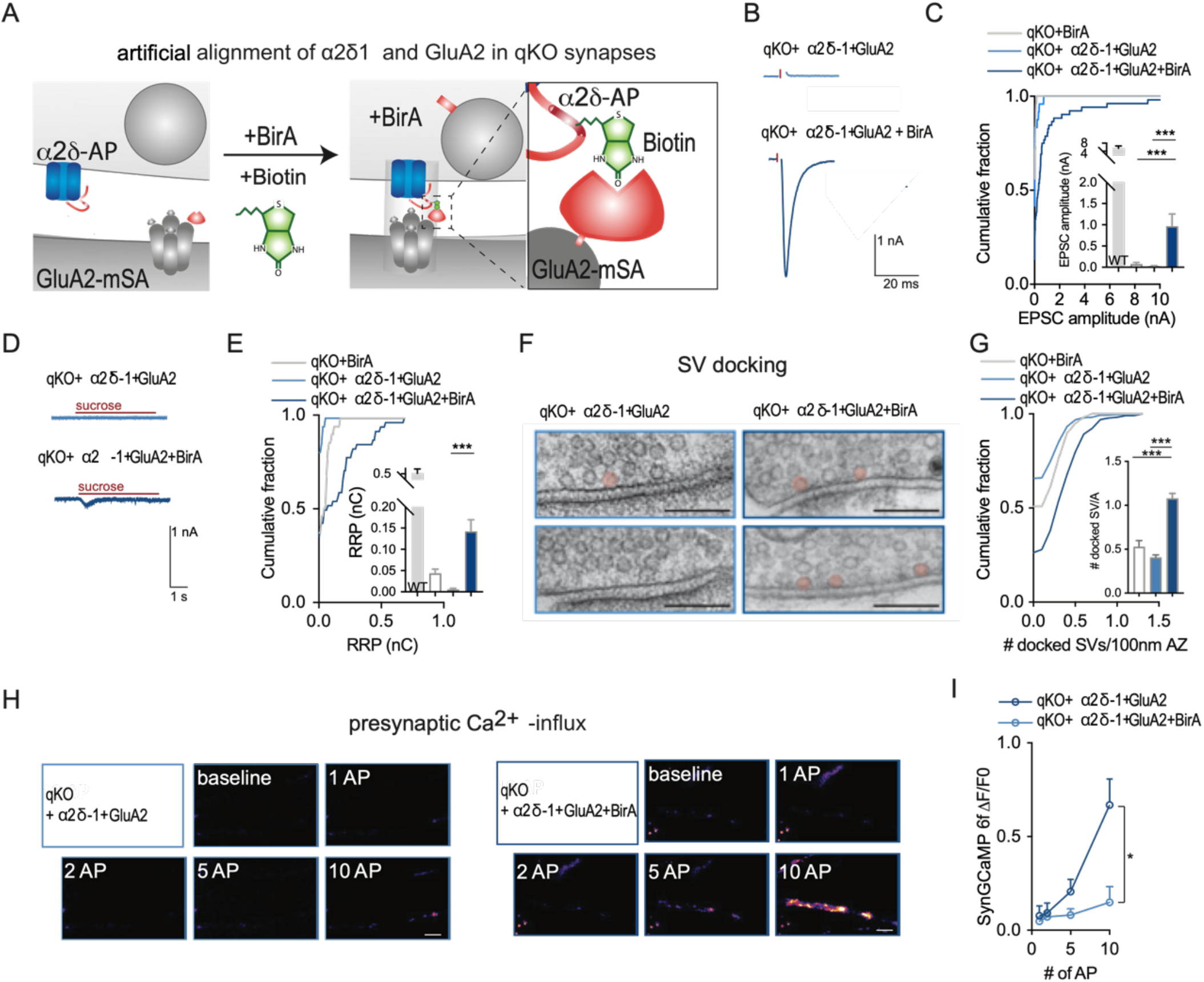
The artificial alignment of α 2δ 1 and GluA2 rescues neurotransmission in RIM/RBP deficient synapses. **(A)** Model of transsynaptic alignment of α2δ1-AP and GluA2-mSA by BirA co-expression in qKO synapses. (**B)** Example traces of excitatory postsynaptic currents (EPSC). (**C)** Excitatory postsynaptic current (EPSC) amplitudes. (**D)** Example traces of synaptic responses to a 5 s application of hypertonic sucrose. (**E)** Readily releasable pool (RRP) charge. (**F)** Representative electron micrographs of cryofixed primary hippocampal neurons from qKO mice. (**G)** Absolute number of docked SV and docked SV per 100 nm AZ. (**H)** Example images of SynGCaMP6f fluorescence in qKO neurons, obtained during trains of AP stimulation. Scale bars 5 µm (**I)** Fluorescence changes (ΔF/F) upon single AP and APs trains. Data are individual values and means ± SEM. Statistical significance for C, E, and G was assessed by Kruskal-Wallis test and I by Two-way ANOVA. *p < 0.05, **p < 0.01, ***p < 0.001.

## Discussion

We developed a novel live labeling EM based protein technique and utilized this technique to quantify -with single-digit nm resolution-the spatial arrangement of the three transducers for neurotransmission: release competent SVs, voltage-gated Ca^2+^-channels, and postsynaptic AMPA receptors at central hippocampal synapses. Interestingly, we found that Ca^2+^-channels are located at a preferred distance of 20 nm to docked SVs and that action potential induced Ca^2+^-influx through Ca^2+^-channels triggers the fusion of these SVs with a release probability of 50 %. Furthermore, we found that the three transducers form a transsynaptic nanocomplex. By artificially reconstituting a transsynaptic interaction between the Ca^2+^-channel auxiliary subunit α2δ1 and the AMPA subunit GluA2, we provide the first evidence that α2δ1 can serve as a nucleation point in the presynapse to align the transducers for neurotransmission, independent from the presynaptic scaffold. This finding proposes a novel mechanism on how individual synaptic units self-organize to orchestrate neurotransmission.

The finding that pre- and postsynaptic scaffold proteins align in nanocolumns (Tang et al., 2016) strongly suggested a transsynaptic association of the transducers for neurotransmission. Indeed, by utilizing our live-gold-labeling of synaptic proteins, we showed for the first time that the pre- and postsynaptic transducers transsynaptically align at a spatial scale equivalent to that of the SV radius. Using the efficacy of slow intracellular Ca^2+^-buffers and diffusion models, previously published works suggest varying coupling distances at mammalian synapses, e.g., 10-20 nm for basket cells, 30-60 nm for the Calyx of Held, and 65-90 nm for mossy fiber synapses (Bohme et al., 2018). Since the key parameter – the distance of fusion-competent vesicles to Ca^2+^-channels – was lacking, the release probability of SVs was simulated. Simulations established that vesicular release is sensitive to their distance to Ca^2+^-channels in a range of 5-10 nm (Bennett et al., 2000; Scimemi and Diamond, 2012), that SVs close to Ca^2+^-channels have a higher release probability than distant once (Meinrenken et al., 2002; Neher, 2015) and that coupling-distances above 100 nm do not contribute to synaptic release (Nakamura et al., 2015). Our study used functional anatomy at small hippocampal synapses, which enabled us to directly determine the spatial coupling and release function of fusion competent SVs and Ca^2+^-channels. Compared to simulated data from other synapses (Nakamura et al., 2015; Rebola et al., 2019), the estimated release probability for small hippocampal synapses is rather high, but detailed modeling and consideration of the fraction and mobility of fusion competent SVs that are not associated with Ca^2+^-channels will likely provide a better understanding how overall release properties and short-term plasticity characteristics are defined by the anatomical state of the synapse.

Deploying our functional anatomy approach, it is now possible to also measure fusion probability as a function of Ca^2+^-channel distance for other synapse types, such as the hippocampal mossy fiber bouton, and compare the physical arrangement of synaptic transducers with the resulting function in synaptic transmission. Comparing the measured distance between docked SVs and Ca^2+^-channels in synapse types with different release probabilities and short-term plasticity characteristics will address whether diversity in coupling distances indeed contributes to varying release efficacies, thus uncovering an underlying mechanism for synaptic diversity. These approaches will not only teach us about the function of individual synapses, but they will also help to add significant new understanding for computation in neural networks when the approach is applied to large scale connectome analysis, as individual connections may be decoded for their release probability.

Our live-gold-labeling technique, paired with high-pressure-freezing EM, revealed the spatial arrangement between membranous structures and pre- and postsynaptic proteins. A previous study, using super-resolution light microscopy, demonstrated the transsynaptic nanocolumn arrangement of synaptic scaffold proteins (Tang et al., 2016). However, visualization of the SV, a crucial component in the neurotransmitter release process, was lacking. By combining EM with efficient protein labeling, we have proven that docked SVs align with Ca^2+^-channels and AMPA receptors. Strikingly, the adhesion molecules neurexin1 and neuroligin1 are not associated with docked SVs and are primarily found at the active zone’s edge, rendering them an unlikely candidate for stabilizing the transducer complex.

What are possible candidates for stabilizing the transducers’ complex, and what drives its assembly? To gain mechanistic insight, we made use of synapses lacking the presynaptic scaffold. The deletion of RIM and RBP results in synaptic loss and disruption of the transducers, shown by a substantial reduction in gold particles labeling Ca^2+^-channels and AMPA receptors. The disruption of the transducers’ alignment results in major synaptic changes, including the loss of docked SVs and neurotransmission. Strikingly, we found that the stabilization of the Ca^2+^-channel subunit α2δ1 at the presynapse in scaffold lacking synapses restores primary presynaptic functions such as the rescue of SV docking and priming, Ca^2+^-influx, and Ca^2+^-triggered release, suggesting that the α2δ1 subunit, rather than the α-subunit (Held et al., 2020), plays a key organizing role in the synapse. These results imply that the Ca^2+^-channel α-subunit that mediates the voltage-dependent Ca^2+^-influx is stabilized by α2δ1. Therefore, the RIM and RBP scaffold that interacts intracellularly with the α-subunit may perform fine-tuning of the voltage-dependent Ca^2+^-channel localization rather than the initial stabilization. So far, we still do not understand the molecular mechanism that stabilizes α2δ at the synapse. The most noteworthy candidates are cis or trans-interacting adhesion molecules, which orchestrate pre- and postsynaptic scaffolds to form nanocolumns (Chen et al., 2017; Sudhof, 2017).

How critical is the presynaptic scaffold in the organization of synapses? While its loss causes severe impairments in synaptic structure and function, synaptic contacts still form, SVs accumulate, and the postsynapse differentiates to some degree (Acuna et al., 2016; Brockmann et al., 2020). Our finding that the artificial alignment of α2δ1 and AMPA receptors suffices to restore synaptic transmission in the absence of the presynaptic scaffold not only suggests that the scaffold is not essential for neurotransmission, it also places α2δ1 as a key component of the presynaptic module that governs presynaptic release. However, the presynaptic scaffold, especially RIM and RBP2, has been proven to fine-tune the coupling distance between Ca^2+^-channels and docked SVs. Therefore, the coupling distances of 20 nm between Ca^2+^-channels and docked SVs and their high release probability of 50% that we revealed for hippocampal synapses are likely mediated by the unique organization of the presynaptic scaffold in that synapse. All in all, our results suggest a molecular model in which a synapse contains autonomous building blocks that govern synaptic structure and -function and a presynaptic scaffold modulating that fine-tunes vesicular release probability.

## Supporting information

Statistics

## Acknowledgements

This research was supported by an Einstein BIH Visiting Fellow grant (EVFBOJ2017-369) and German Research Council grants (RO 1296/8-1; Reinhardt Koselleck project; CRG958A5; C.R.). We would like to thank Melissa Herman for valuable comments on the manuscript. We thank Diana Steinkampf for data analysis. We are grateful to Berit Söhl-Kielczynski, Bettina Brokowski, Katja Pötschke, Heike Lerch, Rike Dannenberg and Sabine Lenz for technical assistance. We also thank for the services of the Charité viral core facility for virus production and the electron microscopy core facility for technical support. We would also like to thank Jan Schmoranzer and the Charité AMBIO facility for the training and use of the STORM setup.

## Author contributions

M.M.B. performed electron microscopy and electrophysiological experiments

E.T. performed super-resolution microscopy experiments

M.M.B., T.C. S., and C.R. designed experiments

A.T.G., A.M.W. performed computational analysis

T.T. designed lentiviral constructs

M.M.B. and C.R. wrote, and A.M.W. edited the manuscript

## Material and Methods

### Animals maintenance and mouse lines

All mouse experiments were in accordance with the regulation of the animal welfare committee of the Charité Universitätsmedizin and the Landesamt für Gesundheit und Soziales Berlin under license number T0220/09. The qKO mouse line contains floxed RIM1, RIM2, RBP1, and RBP2 genes, as published previously(Acuna et al., 2016). The gene deletion is induced by lentiviral transduction of cre-recombinase. Wildtype neurons were obtained from C57Bl/6 mice.

### Primary hippocampal cultures

Hippocampal neurons were dissected from postnatal day (P) 0-2 mice of either sex, as described previously (Arancillo et al., 2013). Briefly, hippocampi were dissected, and neurons dissociated by an enzymatic treatment using 25 units per ml of papain for 45 min at 37 °C. For electron microscopy experiments, 100 × 10^3^ neurons/well (22 mm diameter) were plated on astrocyte feeder layers as high-density culture (Chang et al., 2018). Low-density cultures of 3 × 10^3^ neurons/well were seeded on astrocyte micro-islands (35 mm diameter) for autaptic cultures (Arancillo et al., 2013). For dSTORM experiments, mice hippocampal neurons were plated on 25 mm 1.5H glass coverslips (Carl Roth) following an adapted Banker culture protocol (Kaech and Banker, 2006). Astrocyte feeder layers were prepared 1-2 weeks before neuronal seeding, as described previously (Arancillo et al., 2013). After plating, neurons were incubated in Neurobasal-A media medium supplemented with 50 μg/ml streptomycin and 50 IU/ml penicillin at 37 °C, before electrophysiological, imaging, or electron-microscopy experiments were performed at DIV 14-20.

### Lentiviral constructs

All lentiviral constructs were generated through the Gibson assembly method (NEB) with the corresponding cDNAs and with a human synapsin-1 promoter-driven lentiviral shuttle vector (f(syn), based on FUGW (Lois et al., 2002)) that could contain either nuclear-localized (NLS) GFP or RFP that was fused C-terminally to a self-cleaving P2A peptide (Kim et al., 2011) to allow polycistronic translation.

To perform Ni-NTA-Nanogold labeling for electron microscopy, the sequence of 10 Histidine residues (His) was incorporated in the corresponding cDNAs. For the postsynaptic AMPA receptor GluA2, SEP-GluA2 (Addgene #24001) was used by exchanging the pHluorin cDNA with His and Gibson assembling into f(syn)NLS-GFP-P2A (BL-984). Postsynaptic mouse neuroligin-1 was PCR amplified from Addgene #15260. Here, the His sequence was fused after the N-terminal signaling peptide of neuroligin, and the resulting neuroligin-His was assembled in the f(syn)NLS-RFP-P2A shuttle vector (BL-1045). Presynaptic rat neurexin1a was obtained from Addgene #58266. The His-sequence was also fused after the N terminal signaling peptide. neurexin-1-His was cloned in the shuttle vector f(syn)NLS-GFP-P2A (BL-1047). Presynaptic Cacna2d1 was PCR amplified from mouse brain cDNA P0. After sequence verification, the His-tag was inserted after the signaling peptide sequence by Gibson assembly and fused in the empty shuttle vector to form f(syn)Cacna2d1-His (BL-744).

For super-resolution microscopy, the cDNA of mEOS3.2 (from Addgene #54696) was fused within the cDNA of (i) GluA1 (Addgene #24000) and of (ii) GluA2 (Addgene #24001) by replacing the corresponding pHluorin cDNA within these Addgene clones. GluA1-mEOS3.2 and GluA2-mEOS3.2 were subcloned into the f(syn)NLS-GFP-P2A shuttle vector (BL-1038, BL-1054). From the same Addgene plasmid #57461 Homer1 cDNA was PCR amplified from the Addgene plasmid #57461 and fused at its C-terminus with tdTomato into the shuttle vector f(syn) to produce f(syn)Homer1-tdTomato (BL-1034). The cDNA of presynaptic mouse synaptophysin (Sampathkumar et al., 2016) was fused at its C-terminus with tdTomato into the shuttle vector f(syn) to form f(syn)Synaptopysin-tdTomato (BL-1039).

In order to perform site-specific biotinylation and streptavidin interaction studies, bacterial biotin ligase BirA-ER from Addgene #20856 was subcloned into a CFP-P2A expressing shuttle vector to form f(syn)CFP-P2A-BirA-ER. A 15 amino acid biotin acceptor peptide (AP) (Howarth et al., 2005), was inserted in the Cacna2d1 cDNA by exchanging the His-tag and subsequent assembling in the NLS-GFP-P2A containing shuttle vector to form f(syn)NLS-GFP-P2A-Cacna2d1-AP. As counterpart, the sequence of monomeric streptavidin (mSA) was amplified from Addgene #39860, fused after to the GluA2 sequence (Addgene clone #24001, GluA2-pH) by replacing the pHluorin sequence. Subsequently, this fragment was assembled into an NLS-RFP-P2A containing shuttle vector to form f(syn)NLS-RFP-P2A-GluA2-mSA.

### Ni-NTA gold labelling

For the Nickel Nitrilotriacetic acid (Ni-NTA) labeling, primary hippocampal neurons were grown on sapphire disks for 14 days. Before high-pressure freezing, neurons were incubated in 10 nM 5 nm Ni-NTA-Nanogold (Nanoprobes) (Chang et al., 2013) for 15-20 min in the CO^2^ incubator at 37°C. To remove unbound gold particles, cells were washed ten times in Base^+^ solution at RT that contained the following (in mM): 140 NaCl, 2.4 KCl, 10 HEPES (Merck, NJ, USA), 10 glucose (Carl Roth, Karlsruhe, Germany), 2 CaCl_2_(Sigma-Aldrich, St. Louis, USA), and 4 MgCl_2_ (Carl Roth) (∼300mOsm; pH7.4). Primary neurons were high-pressure fixed ∼2 min after finishing the washing steps.

### High-pressure freezing with electrical stimulation

For high-pressure freezing experiments, hippocampal neurons were plated on 6 mm sapphire disks. Neurons were high-pressure fixed (2100 bar, ICE, Leica Microsystems) between DIV 14-16 in extracellular solution containing the following (in mM): 140 NaCl, 2.4 KCl, 10 HEPES, 10 glucose, 2 CaCl_2_, and 4 MgCl_2_ (∼300mOsm; pH7.4). For the gold labeling, neurons were incubated with the Ni-NTA 15-20 min before freezing. All experiments without stimulation were performed at room temperature. For electrical stimulation 3 μM NBQX and 30 μM bicuculine were added to the extracellular. To induce vesicle fusion, the ICE applies an electrical field for 1 ms, and cells were frozen 5 ms after stimulation at 37 °C.

After freezing, samples were transferred from liquid nitrogen to cryovials containing the freeze substituent (1 % glutaraldehyde, 1 % osmium tetroxide, and 1 % H_2_O in anhydrous acetone). The freeze-substitution was performed in an automated freeze-substitution (Leica EM AFS2) with the following protocol: −90°C for 5-7h, 5°C per hour to −20°C. 12 hours at −20°C, and 10°C per hour to 20°C. Following en bloc staining with 0.1% uranyl acetate for 1 hour, the samples were epon embedded and cured for 48 hours at 60°C. Afterward, 40 nm sections were cut using a microtome (Leica UCT). The sections were collected on formvar-coated single-slot grids and stained before imaging with 2.5% uranyl acetate and lead citrate. Pictures were collected blindly in an FEI Tecnai G20 TEM operating at 200 keV and digital images taken with a Veleta 2 K x2 K CCD camera (Olympus). For Tomograms, 200 nm sections were obtained and imaged with an FEI Tecnai G20. Tomograms were obtained from synapses with at least 1 gold particle detected and reconstructed and analyzed using IMOD (University of Colorado, Boulder; http://bio3d.colorado.edu/imod/).

For all experiments, the images were obtained from at least 3 independent hippocampal cultures. Synapses for imaging were chosen by the presence of a prominent PSD, presynaptic vesicles, and a visible synaptic cleft in low magnification. The images were collected and scored blind by using an analysis program for ImageJ and Scala. Active Zones were defined as the membrane stretch directly juxtaposed to the postsynaptic density (PSD). Docked SVs were defined as those contacting the active zone plasma membrane.

### 2D electron microscopy data analysis

#### Synaptic Parameters analyzed

Morphological analysis of EM images performed manually in ImageJ. Micrographs were blinded for the experimenter to prevent biased analysis. In Fiji, the post-synaptic density (PSD) was defined as the electron-dense structure at the postsynapse and the active zone as the membrane stretch opposed to the PSD at the presynaptic side. Vesicles were defined as docked when the vesicular membrane directly contacted the presynaptic plasma membrane. Gold particles were marked in the synaptic cleft when found in the confines of the active zone. After detecting all relevant structures, localization parameters were exported as a text file.

#### Distance calculation

To determine the distances between gold particles and docked SV, coordinates of all structures were loaded from each text file into a self-written analysis program. As the image files contain only points, we transformed the data in the following way in order to be able to compute distances: the AZ and the PSD were approximated using a 3^rd^-degree polynomial and fitted with the least error squares. Subsequently, the polynomial was divided into straight line segments by computing the respective values on the highest resolution available from the data (once per pixel). We projected the docked SVs and the gold particles onto the line segments by using the point which has the minimal distance across all line segments.

We then calculated the following distances on the line segments: 1) The length of the PSD. 2) The position of the gold particle along the AZ length. 3) The number of gold particles in the confines of the synaptic cleft. 4) Number of docked SV. 5) Location of docked SV along the AZ length. 6) Distance between docked SVs. 7) Distance between each gold particle and the nearest docked SV. 8) Distance of each gold particle to all docked SVs.

#### Randomized placement of SVs and gold particles for 2D data

To analyze gold and docked SVs distribution, measured data were compared to simulations of randomly equally distributed SVs and gold particles. For randomized conditions, the measured number of SVs and gold particles were randomly placed on the given AZ length.

#### Residual value calculation

To obtain the residual value (%), frequency distribution histograms (bin 20 nm) for were utilized and calculated as the percent of increase distance of docked SVs to Ca^2+^-channels.

### Image analysis tomograms

#### Randomized placement of SVs and gold particles

If a guided process determined SV placement in relation to Ca^2+^-channels, then the distribution of distances between docked SVs and Ca^2+^ channels should differ from a random uniform distribution. To test this hypothesis, we first quantified the shortest distance of each immunogold particle (“au”, as observed in 3D EM tomographs) to the closest observed docked SV, yielding one distance value per observed au (n=19 in 10 AZ tomographs). We then simulated randomly placed positions of docked SVs and au by randomly drawing AZ coordinates for their uniform placement, as described below. All following procedures were performed in MATLAB (R2016b). The custom scripts written for this analysis are available upon request.

#### Data handling and distance calculations

AZ coordinates were loaded from .xlsx-files containing one AZ per sheet (in this case 10 sheets for 10 analyzed AZs), with data being described by 5 columns. In the first column, all AZ coordinates were classified as “1”, all docked SV positions were classified as “2”, and all au positions were classified as “3”. The second column contained information about the tomograph slice identity but was not used in this analysis. The third to fifth column contained x, y and z coordinates of the respective AZ, SV or au position. Each sheet was then loaded into a MATLAB cell array using *xlsread* and the 3^rd^ to 5^th^ columns were converted to a 3-column matrix using *cell2mat*. Then, going through each AZ individually, the following information was stored: the index of AZ coordinates (“1”), the index of docked SV positions (“2”), and the index of au positions (“3”), yielding three cell arrays containing either AZ, docked SV or au coordinates. Then, for each docked SV and au, we projected its position onto the AZ surface by finding the AZ x,y,z-coordinate closest to the x,y,z-coordinates of the docked SV or au. The distances between SV/au and AZ coordinates were determined using *pdist2*, which calculates Euclidean distances between given sets of points, and taking the smallest value from this list, as shown in equations (1) and (2).

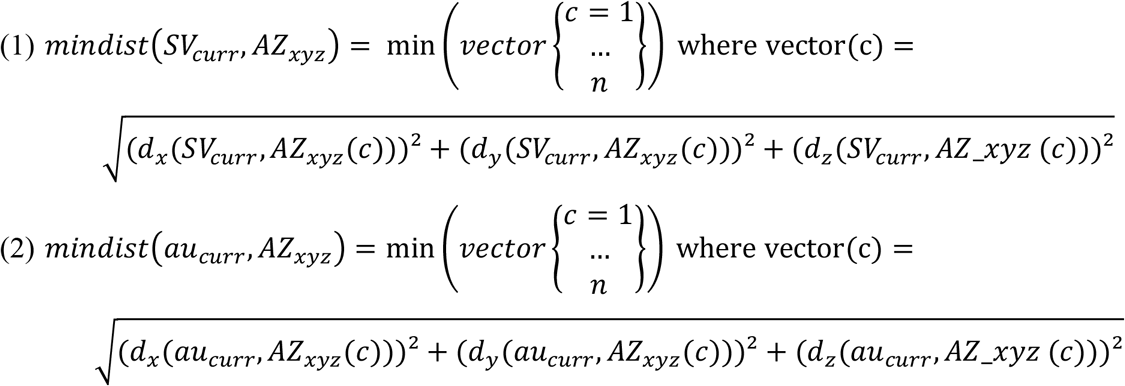

In equations (1) and (2), SV_curr_ and au_curr_ are the docked SV or au being considered, and AZ_xyz_ is the set of all AZ x,y,z-coordinates. For each single x,y,z-coordinate AZ_xyz_(c), d_x_, d_y_ and d_z_ are the difference (or “shift”) between SV or au and the currently considered AZ_xyz_(c) along either spatial dimension (x, y or z)). The counter c increases from 1 to n, where n is the amount of observed AZ coordinates. The x,y,z-coordinate AZ_xyz_ corresponding to the shortest distance is then taken as the projected position of the docked SV or au.

Having determined the projected coordinates of all au particles and docked SVs, we calculated the distances between each au particle and the closest docked SV, resulting in as many distance measurements per AZ as there were observed au particles (n = 19). Because a possible curvature of the AZ surface is not easily quantified, exact distance measurement along the 3D surface coordinates poses a non-trivial problem. However, AZs were generally rather flat in shape. We therefore approximated the actual surface distance by taking the shortest Euclidean distance in space, analogous to equations (1) and (2) and as schematically depicted in Figure S4. This was done for both the experimental and simulated data.

#### Simulated random placement and bootstrapping

To generate a distribution with randomly placed components, we proceeded as follows. For each observed AZ, we placed as many docked SVs and au particles as had been observed in that AZ in EM tomography. Placement happened by choosing a random position of their projection on the AZ surface (using the MATLAB function *randi*) to randomly determine an integer number that corresponded to a position in the list of all observed AZ x,y,z-coordinates from a uniform distribution. We further excluded unrealistic cases by excluding placements where SVs were positioned less than 40 nm apart (because docked SVs cannot physically overlap, assuming an SV diameter of 40 nm) by randomly redrawing positions until all SVs were sufficiently far apart. This procedure was performed analogously for randomly placed au particles, but no minimum distance between au particles, or between au particles and SVs, was enforced.

Next, we wanted to compare the distribution of observed minimal distances between au and docked SVs with the distribution of randomly uniformly placed au and docked SVs. To account for effects of variability within the experiment on the outcome, we furthermore performed bootstrapping (using the MATLAB function *bootstrp*) on both experimental and simulated datasets. For the experimental data, this was done from the original dataset of 10 AZs by drawing 100 subsets from the 10 AZs and generating an average histogram of the derived distances in 20 nm bins. This was done by first binning the distances in 20 nm bins and then averaging over all 100 bootstrapping runs. Calculating the cumulative sum over distance bins resulted in the plot shown in Figure 1G (α2δ-His).

The random drawing of simulated random positions in 10 AZs was then performed 100 times, using the same bootstrapped AZs as in the above procedure, where au particles and SVs were randomly uniformly placed in each bootstrap run and the observed positions quantified. For each random drawing of positions in 10 AZs, we binned the observed minimal distances between au and docked SVs. After repeating this 100 times, we then generated the average histogram we could compare to the distribution of observed distances. The cumulative distribution of this random drawing is shown in shown in Figure 1G (simulated).

### Fitting the minimal distance distribution for EM tomograms

We utilized an integrated Rayleigh function that describes the distribution of distances between docked SVs and gold-labelled α2δ (Figure 1H) with a single parameter, *σ*, similar to analysis done previously at the NMJ active zone of L3 stage drosophila^39^. A maximum likelihood estimation using the Matlab function *mle* was used to estimate *σ*. For comparison, *σ* was 3.8 times larger at the *Drosophila* NMJ (using light microscopy; *σ* =76.52 nm^39^) than our estimates at the central mammalian synapse, arguing for an overall shorter coupling between voltage gated calcium channels and docked SVs.

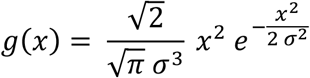

### Calculation of the vesicular release probability as a function of distances to α2δ gold label

The profiles depicting the distance-dependence of the vesicular release probability (pV_r_, Figure 2D) were calculated by comparing the measured minimal distances between gold particles and SVs without stimulation to the ones observed 5 ms after stimulation that triggered SV fusion. For the quantification depicted in Figure 2D, the analysis was performed on EM profiles containing a single gold particle only. This simplifies the assignment of SVs to the respective particles and alleviates the concern that the number of gold particles may influence the observation frequency of distances (as the average number of gold particles observed in the two groups were different). The observation frequency of distances between gold particles and docked SVs were quantified in the control dataset in 6 bins whose sizes were chosen such that a same/similar number of observations fell into those bins. The probability density of observations was then calculated by dividing the number of observations per bin by the respective bin-width. The same bins were then used to calculate the probability density of distance observations between gold particles and docked SVs in the profiles obtained 5 ms after stimulation. In the latter the overall number of observations was reduced, consistent with the loss/triggered membrane fusion of docked SVs following the stimulus. To investigate the variability within the datasets, bootstrapping of the observed distances was performed in both cases with 10000 bootstrap runs by drawing samples of distances using the second output of the Matlab function *bootstrap*. The respective numbers of observations were then counted using the using the Matlab (release Matlab2017b, Mathworks, MA, USA) function *histcounts* in the 6 bins and this number divided by the respective bin width to obtain the observation density. This number was furthermore divided by the number of profiles analyzed in the respective groups (57/196 profiles in the case of the control dataset and 62/204 profiles following stimulation for the analysis of profiles containing a single /at least one gold particle). The vesicular release probability (pV_r_) for all 10000 bootstrap runs in each bin was then calculated by subtracting the probability density of observed distances after stimulation from the ones without stimulation and then dividing by the probability density of observations without stimulation. In some cases, also depending on the bootstrap sample, negative values were calculated when a higher observation density is observed in the group following stimulation than in the control group. Because negative probabilities are not defined, the pV_r_ value was then forced to zero (the smallest defined value). This typically occurred for larger distances, where overall fewer docked SVs were observed in the control condition. Figure 2D depicts the mean pV_r_ profile plotted against the bin center values plus and minus the standard deviation from the bootstrapped data. The mean pV_r_ profile was fit (using the Matlab function *fit*) with a single exponential decay function

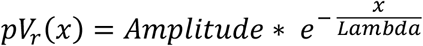

where *Amplitude* denotes the (maximal) pV_r_ value at the position of the gold particle, *x* denotes distance from the gold particle and *Lambda* is a length constant. The values of the parameters Amplitude and Lambda were found by optimization (from starting values 1 and 30 nm) by the fitting routine.

### Electrophysiology

Whole-cell patch-clamp recordings were performed on autaptic cultures at room temperature at days in vitro 15-20. Synaptic currents were recorded using a Multiclamp 700B amplifier (Axon Instruments) controlled by Clampex 9.2 software (Molecular Devices). A fast perfusion system (SF-77B; Warner Instruments) continuously perfused the neurons with the extracellular solution contained the following (in mM): 140 NaCl, 2.4 KCl, 10 HEPES (Merck, NJ, USA), 10 glucose (Carl Roth, Karlsruhe, Germany), 2 CaCl_2_ (Sigma-Aldrich, St. Louis, USA), and 4 MgCl_2_ (Carl Roth) (∼300mOsm; pH7.4). Somatic whole cell recordings were obtained using borosilicate glass pipettes, yielding a tip resistance of 2-3.5 MΩ and filled with the following internal solution (in mM): 136 KCl, 17.8 HEPES, 1 EGTA, 4.6 MgCl_2_, 4 Na_2_ATP, 0.3 Na2GTP, 12 creatine phosphate, and 50 U/ml phosphocreatine kinase (∼300 mOsm; pH7.4). Membrane capacitance and series resistance were compensated by 70% und data filtered by low-pass Bessel filter at 3 kHz and sampled at 10 kHz using an Axon Digidata 1322A digitizer (Molecular Devices).

Neurons were held at −70 mV, and action potentials triggered by a 2 ms depolarization to 0 mV to measure EPSCs (excitatory postsynaptic currents). Afterward, EPSCs were induced in the presence of the competitive AMPA receptor antagonist NBQX (3 μM, Tocris Bioscience). Paired-Pulse stimulation for excitatory autapses was assessed by the induction of 2 action potentials with an interstimulus interval of 25 ms. Spontaneous events (mEPSCs) were detected for 60 s, and false positive mEPSC events obtained in NBQX were subtracted to calculate the frequency of spontaneous events. The readily-releasable pool (RRP) was determined by 5 s application of external solution made hypertonic by adding 500 mM and integrating the transient inward response component (Rosenmund and Stevens, 1996). The P_v_r was calculated by dividing the average charge of the EPSC by the RRP charge. Data were analyzed using AxoGraph software.

### Sample preparation and immuno-labeling for dSTORM

For dSTORM imaging of Ca^2+^-channels, neuronal cultures were fixed by incubating cells with a solution of 4% PFA and 4% sucrose for 10 minutes at room temperature before incubation with primary antibody (Abcam, ab6556). In another set of experiments, to ensure labeling of surface expressed GluA1-SEP or GluA2-SEP receptors prior to fixation with 4% PFA and 4% sucrose, primary cultures were live labeled by incubating them with rabbit anti-GFP antibody (Abcam, ab6556) diluted in Ringer solution at 1:2000 for 10 minutes at 37°C. After fixation, PFA autofluorescence was quenched with NH_4_Cl (50 mM) for 5 minutes. Next, cells were permeabilized with 0.3% Triton-X100 for 5 minutes. After 3 washes of each 5 minutes with PBS, nonspecific staining was blocked by incubating coverslips in 2 % BSA solution for 45 minutes at room temperature. Primary antibodies were revealed with goat anti-rabbit Alexa 647 (dSTORM) secondary antibodies (1:1000, ThermoFisher). Incubation with secondary antibody was followed by a second fixation step with 2% PFA and 2% sucrose for 10 minutes, and by an extra step of PFA quenching with the incubation of NH_4_Cl for 5 minutes. Fixed neuronal cultures were kept for up to 2 weeks in PBS prior to imaging.

### Imaging buffer and sample mounting

The STORM imaging buffer was prepared with 10 mM cysteamine (Sigma-Aldrich), 10% w/v D-glucose (Sigma-Aldrich), 0.5 mg/ml glucose oxidase (Sigma-Aldrich), and 40 μg/ml catalase (Sigma-Aldrich) in Tris buffer. 100 µl imaging buffer was dropped onto a spherical void (Carl Roth) before the coverslip containing sample was inverted over the chamber and gently pressed down into a central position. Excess imaging buffer was removed, and the coverslip was sealed in place using picodent twinsil® to minimize oxygen exchange during acquisition.

### direct STochastic Optical Reconstruction Microscopy (dSTORM)

dSTORM experiments were done on fixed immunolabeled neurons. dSTORM imaging experiments were conducted on a Nikon Eclipse-Ti inverted microscope using a Nikon 100x 1.49 NA oil immersion TIRF objective. The perfect focus system (PFS) was used to minimize focal drift during long acquisitions. Excitation light was provided by a fiber-coupled laser launch (405 nm, 488 nm, 561 nm, and 647 nm). The fluorescent signal was collected with an EMCCD camera (Andor iXon Ultra, Oxford Instruments, USA). Image acquisition and control of the microscope were driven by NIS-Elements imaging software (Nikon, USA). Each image stack contained typically 30,000-80,000 frames. Selected ROI (region of interest) had a dimension of 256×256 pixels (one pixel edge was 107 nm). The power of the 647 nm laser was kept constant at 100% during the entire acquisition.

### Cluster analysis

Localization of Alexa-647 signals and Single-Molecule Localization (SML) image reconstruction with a pixel size of 21 nm was performed using the ImageJ plugin Thunderstorm (DOI: <u>10.1093/bioinformatics/btu202</u>). GluA receptor and Ca^2+^-channel receptor localizations were extracted from super-resolved images corrected for lateral drift. GluA and Ca^2+^-channel clusters were then identified using SR-Tesseler software (doi: 10.1038/nmeth.3579). Clusters were extracted in three sequential steps for each image. First, the tessellation of the image creates a Voronoi diagram, which consists of polygons based on the density of localizations. Then a region with a density factor (DF) greater than 2 is defined as an object. Finally, an automatic threshold of normalized density DF = 2 was used to extract clusters of proteins (cluster of level 2) having an enrichment factor higher than the average localization density. For cluster determination, we added 2 extra parameters, specifying a minimum area of 0.002 μm^2^ and a minimum localization number per cluster of 5. The extraction of clusters was performed on the SML image, and the synaptic regions were pre-determined by low-resolution imaging of synaptophysin-tdTomato or Homer-tdTomato according to the particular experiment.

### SynGCaMP6f Imaging

Hippocampal autapses were transduced at DIV 1-2 with lentiviral constructs of synGCamp6f (Brockmann et al., 2020; Grauel et al., 2016). Live imaging at room temperature was performed at an inverted microscope, and samples were continuously perfused with external solution containing NBQX/bicuculline. Neuronal stimulation was achieved by field stimulation (Warner Instruments, CT, USA). Imaging experiments were performed at DIV 15-20 of autapses in response to single stimuli and stimuli trains of 10 Hz. Images were acquired using a 490-nm LED system (pE2; CoolLED) at a 2 Hz sampling rate with 50 ms of exposure time.

### Immunostaining and quantification

Primary hippocampal mass cultures were fixed for 10 min in 4 % paraformaldehyde (PFA), permeabilized with 0.1 % Phosphate Buffered Saline (PBS)-Tween-20 solution and blocked with 5 % normal donkey serum. Primary antibodies were used to immune stain Munc13-1 (sysy 126104) and synaptophysin1 (sysy 101002). Subsequently, secondary antibodies labeled with Alexa Fluor 647 anti-guinea pig IgG, and Alexa Fluor 488 anti-rabbit both in donkey (Jackson ImmunoResearch) serum were applied, respectively. Glass coverslips were mounted on glass slides and images acquired with an Olympus IX81 epifluorescent microscope with MicroMax 1300YHS camera using MetaMorph software (Molecular Device). The analysis was performed offline with ImageJ.

### Statistical Analysis

Statistical tests were performed with Prism 8 (Graphpad). For bar plots data are represented as mean ± SEM and for violin plots data were presented as median (solid lines) and quartiles (dashed lines). First, all data were tested for normality using the D’Agostino & Pearson test. If they did not pass the parametric assumption, for bar graphs the Mann-Whitney U-test for the comparison of two unpaired data sets. For cumulative distribution plots the Kruskal-Wallis test followed by Dunn’s test was performed. In case data passed the parametric assumption, for data sets including two experimental groups we applied the Student’s t-test and for analysis an Ordinary One-way ANOVA followed by Turkey’s test. For detail see Supplementary Table 1.

All experiments in the manuscript are repeated at least three times.

## Data and materials availability

Data and Materials of this study are available from the corresponding authors upon reasonable request.

## Code availability

Custom codes used in this study are available from the corresponding authors upon reasonable request.

---

**Figure S1:**
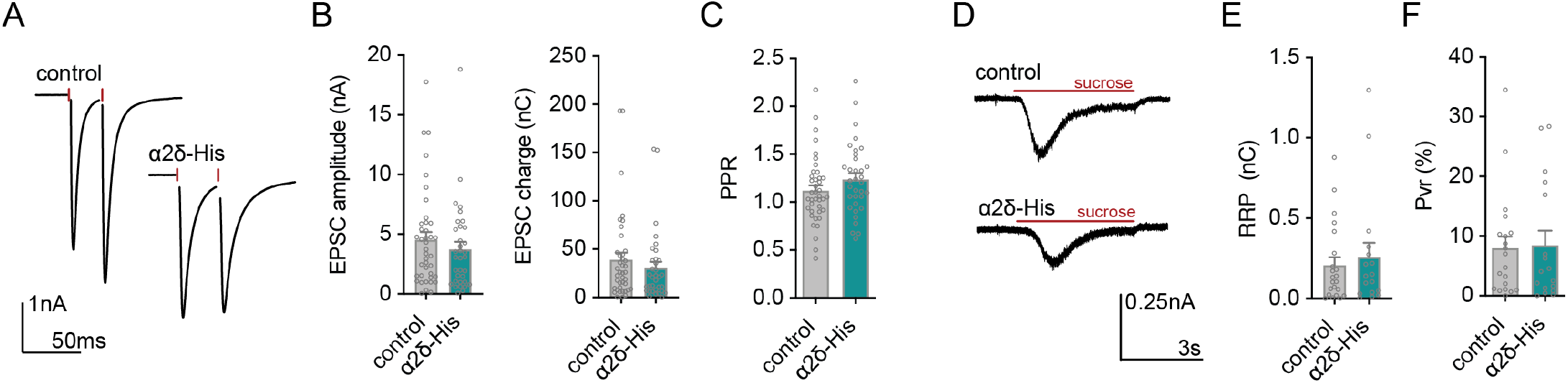
The lentiviral expression of α 2δ 1-His does not impact excitatory synaptic transmission. **(A)** Example traces of two EPSCs evoked by a 2 ms depolarization with an interstimulus interval of 25 ms from control and α2δ1-His expressing hippocampal autapses. (**B)** Summary graphs of excitatory postsynaptic current (EPSC) amplitude, and charge. (**C)** Summary graph of the Paired-Pulse-ratio (PPR) calculated as the fraction of the second EPSC and the first EPSC at an interstimulus interval of 25 ms. (**D)** Example traces of synaptic responses to a 5 s application of hypertonic sucrose (500 mM) solution probing the readily releasable pool (RRP). (**E)** Summary graph of the RRP. **(F)** Summary graph of the vesicular release probability (P_vr_) calculated as the ratio of the EPSC and the RRP charge. Data are individual values and means ± SEM. Statistical significance for B, C, E, and F was assessed by unpaired t-test.

**Figure S2:**
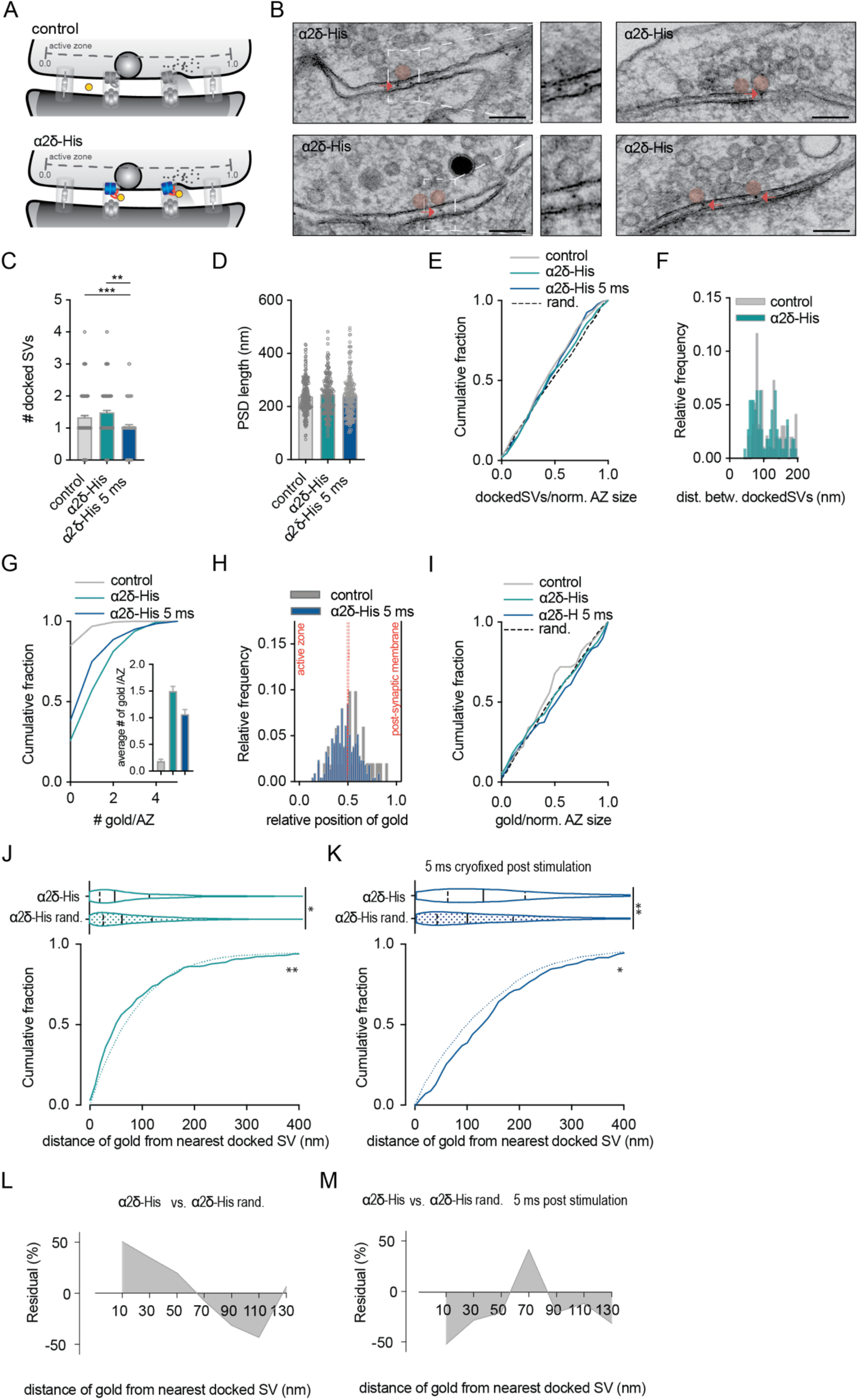
The lentiviral expression of α 2δ 1-His does not impact the ultrastructure of excitatory synapses. **(A)** Scheme of the ultrastructural parameters analyzed in electron micrographs for control (non-transfected) and α2δ1-His expressing neurons. The active zone (AZ) is defined as the membrane stretch opposed to the postsynaptic density (PSD). Synaptic vesicles (SV) are docked when their membrane is directly attached to the AZ membrane. For some analysis, the AZ length is normalized to 1 with the center at 0.5. (**B)** Representative electron micrographs of primary hippocampal neurons expressing the tagged Ca^2+^-channel subunit α2δ1-His. Boxed docked vesicle enlarged together with the respective gold particle. Scale bars, 100 nm. (**C and D)** Summary graphs show the number of docked SVs (c) and the PSD length (d) for control, α2δ1-His expressing neurons, and α2δ1-His expressing neurons cryofixed 5 ms after action potential induction. (**E)** Cumulative probability plot of docked SVs distribution at the normalized AZ. (**F)** Frequency plot of the distance between neighboring docked SVs at the AZ. (**G)** Cumulative probability plot and bar graph of the number of gold particles in the synaptic cleft in control, α2δ1-His expressing neurons without stimulation, and α2δ1-His expressing neurons cryofixed 5 ms after action potential induction. (**H)** Frequency plot of the gold particle position in the synaptic cleft (0 = AZ; 1 = PSD). (**I** Cumulative probability plot of gold particle distribution throughout the normalized AZ length. (**J)** Violin, and cumulative probability plot of the nearest distance of a gold particle to the next docked SV in α2δ1-His expressing neurons compared to random placement of SVs and α2δ1-His. (**K)** Violin, and cumulative probability plot of a gold particle’s shortest distance to the next docked SV in α2δ1-His expressing neurons cryofixed 5 ms after action potential induction compared to random placement of SVs and α2δ1-His. (**L)** Enrichment of a2d1-His localization to docked SVs in 20 nm bins compared to the simulated distribution of random docked SVs and a2d1-His placement. The residual percentage is calculated as the percentage of the measured distances of gold particles to docked SVs to their randomized distribution distances. (**M)** Residual percentage of a2d1-His localization to docked SVs 5 ms after action potential induction in 20 nm bins, measured as the percentage of the measured distances of gold particles to docked SVs to the distances of their randomized distribution. Data for bar graphs are individual values and means ± SEM. Data for violin plots are medians (solid lines) and quartiles (dashed lines). Statistical significance for C, D and G was assessed by Kruskal-Wallis test for the violin plots in J and K by Mann-Whitney test, and for the cumulative distribution plots in E, I, J, and K by the Kolmogorov-Smirnov test. *p < 0.05, *** p < 0.001.

**Figure S3:**
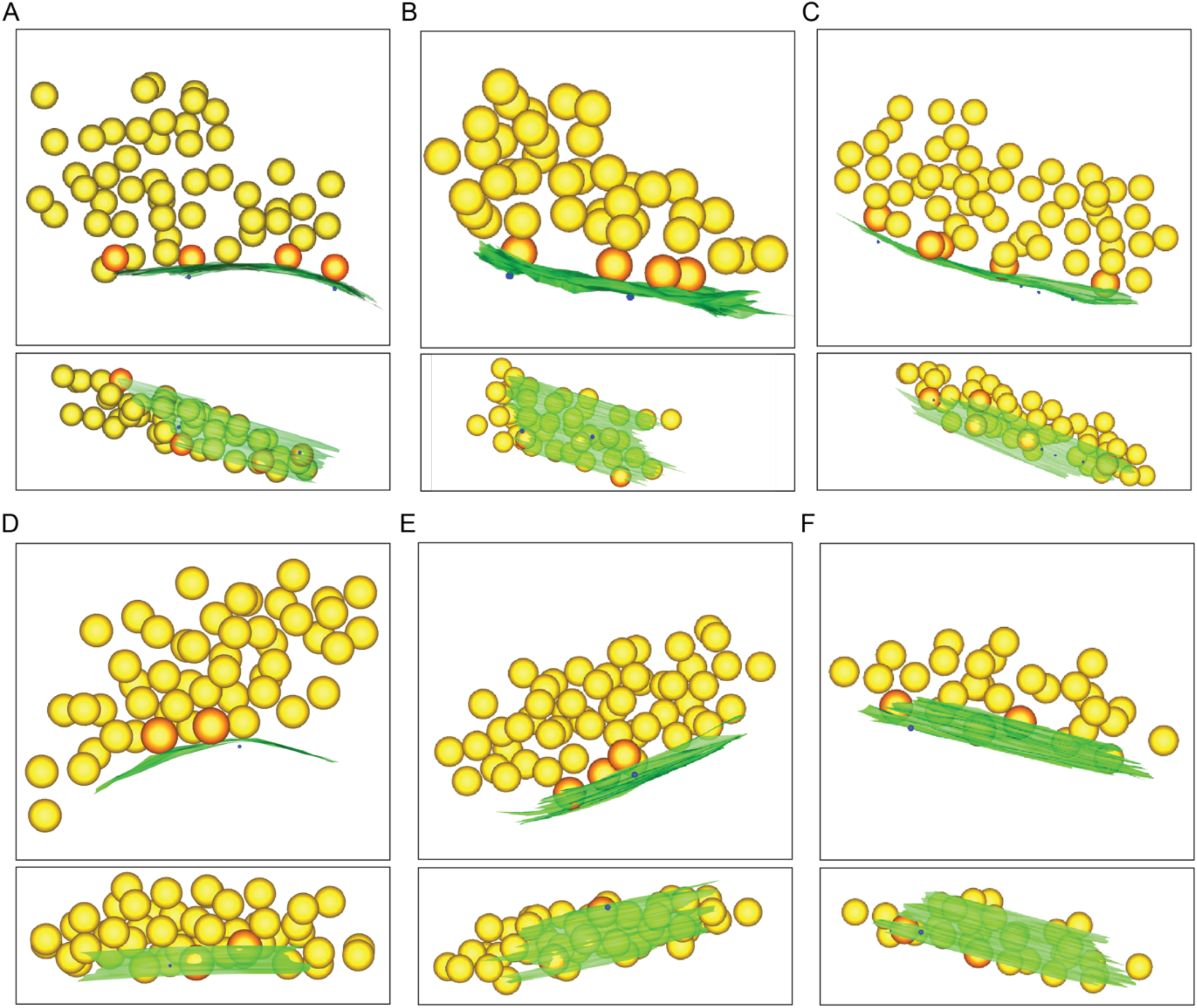
α 2δ 1-His localization in tomograms from cryofixed excitatory hippocampal neurons. **(A-F)** Individual tomograms of synapses from cryo-fixed hippocampal neurons expressing α2δ1-HIS (active zone = green, SVs = yellow, docked SV = orange, gold particle = blue).

**Figure S4:**
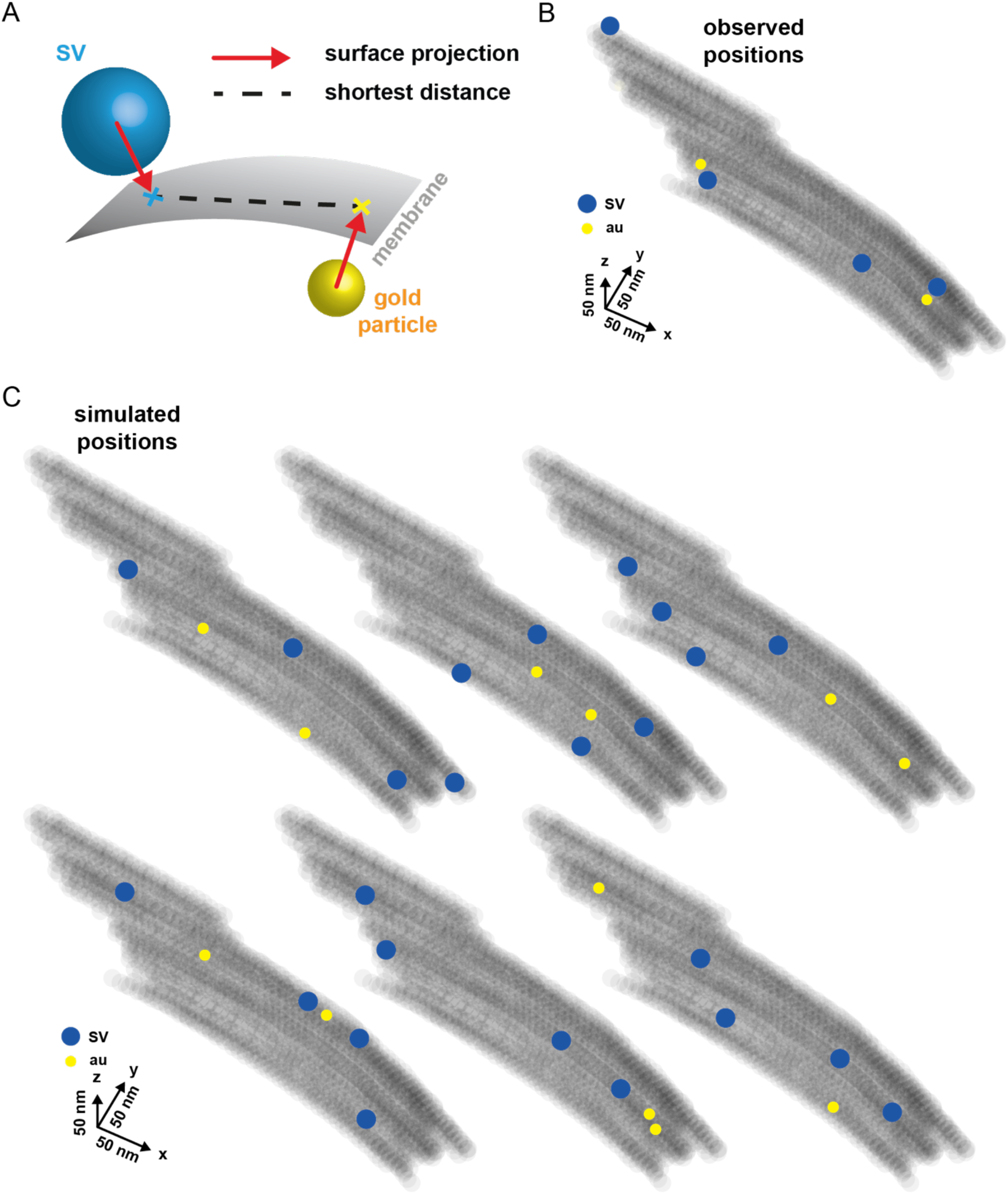
Analysis of α 2δ 1-His localization in tomograms from cryofixed excitatory hippocampal neurons. **(A)** Graphic of the analysis strategy to determine the distance between the gold particle and docked SV from tomogram data. (**B)** Reconstruction of the observed and (**C)** randomized position of gold particles and docked SVs from tomograms.

**Figure S5:**
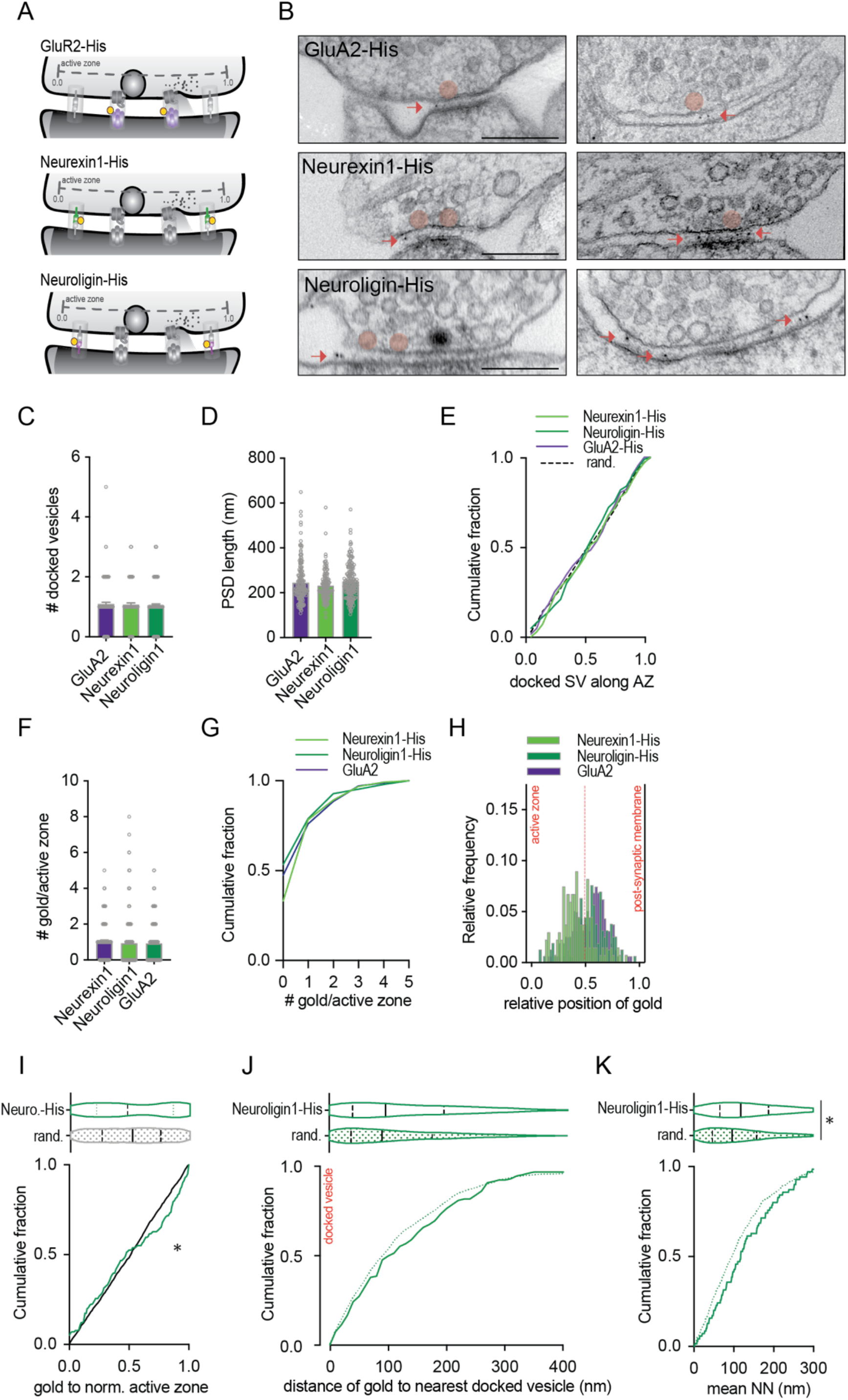
Neuroligin1 does not associate with docked SV. **(A)** Graphic for the ultrastructural parameters analyzed in electron micrographs for GluA2-His, neurexin-His, and neuroligin-His expressing neurons. (**B)** Representative electron micrographs of cryofixed primary hippocampal neurons expressing GluA2-HIS, neurexin1-His, or neuroligin1-His. (**C)** Summary graph of docked SV number per AZ. (**D)** Summary graph of the PSD length. (**E)** Cumulative probability of docked SV distribution at the AZ. (**F)** Summary graph of gold particles per AZ. (**G)** Cumulative probability plot of gold particles per AZ. **h)** Relative frequency of gold particles in the synaptic cleft (0 = active zone membrane; 1 = post-synaptic membrane). (**I)** Violin, and cumulative probability plots of gold the particle distribution at the normalized AZ for neuroligin-His expressing neurons compared to randomized data. (**J)** Violin, and cumulative probability plots of the nearest distance between gold participles and the next docked SV compared to randomized distances. (**K)** Violin, and cumulative probability plots of the measured data and null model data of mean nearest neighbor distance (NND) of gold particles to docked SVs. Data for bar graphs are individual values and means ± SEM. Data for violin plots are medians (solid lines) and quartiles (dashed lines). Statistical significance for C, D, and F was assessed by Kruskal-Wallis test, for I, J, and K by Mann-Whitney test, and for the cumulative distribution plots in I, J, and K by the Kolmogorov-Smirnov test. *p < 0.05.

**Figure S6:**
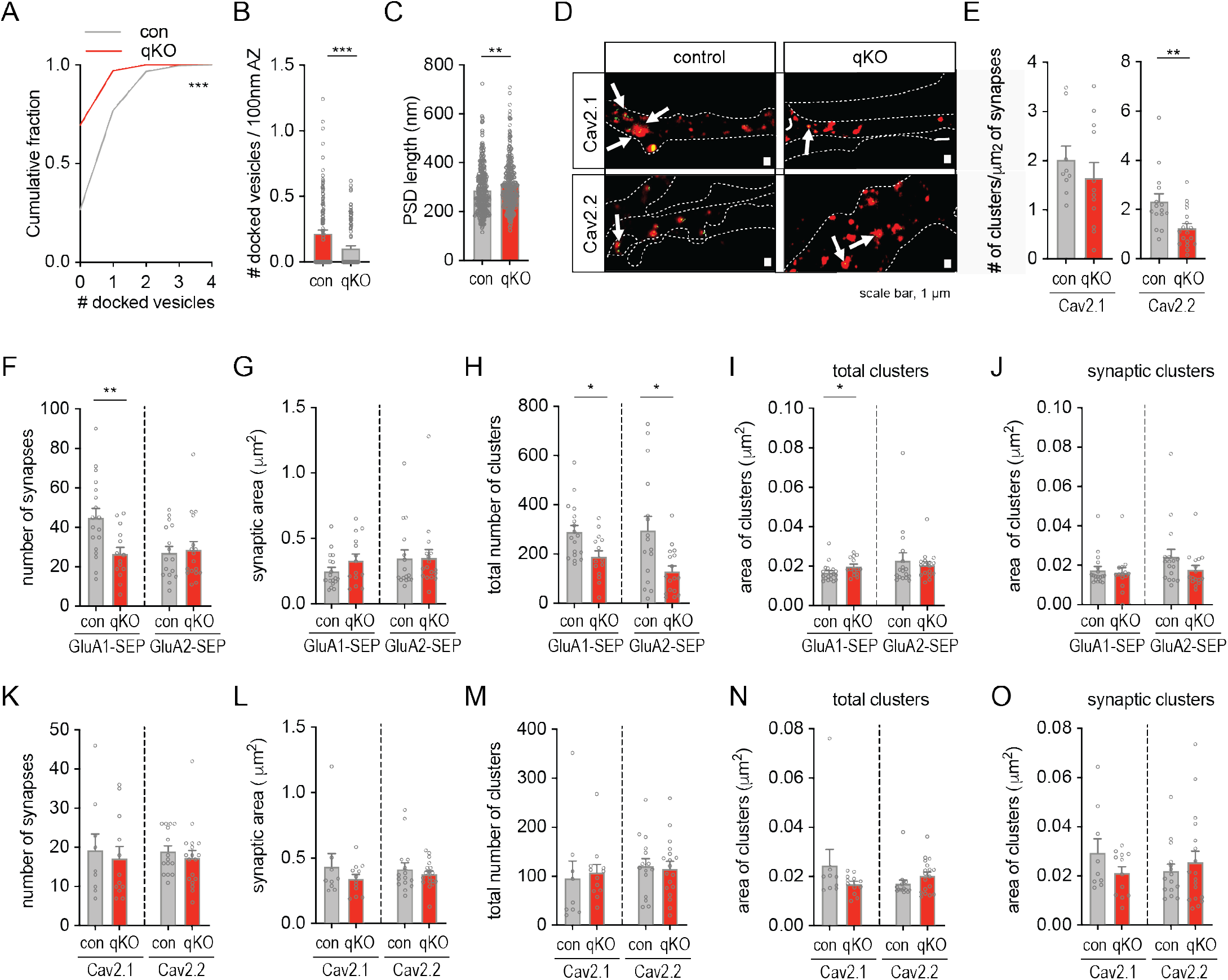
The deletion of RIM/RBP results in a loss of docked SV and a strong reduction in Ca^2+^-channels. **(A-O)** All experiments were conducted in RIM/RBP deficient neurons (qKO) and their respective control. (**A)** Cumulative probability plot of the number of docked SV. (**B)** Summary graph of the number of docked SV per 100nm AZ. (**C)** Summary graph of the PSD length. (**D)** Synapse expressing Cav2.1-GFP or Cav2.2-GFP (green; dSTORM) and Homer (red). Scale bars 1 µm. (**E)** Summary graphs of the number of Cav2.1 and Cav2.2 clusters per synapse area. (**F and K)** Summary graphs of synapse number of the respective AMPA or Ca^2+^-channel subunit. (**G and L)** Summary graphs of the synaptic area of the respective AMPA or Ca^2+^-channel subunit. (**H and M)** Summary graphs of the total number of clusters of the respective AMPA or Ca^2+^-channel subunit. (**I and N)** Summary graphs of the area of all clusters for the respective AMPA or Ca^2+^-channel subunit. (**J and O)** Summary graphs of the area of synaptic clusters for the respective AMPA or Ca^2+^-channel subunit. Data are individual values and means ± SEM. Statistical significance for B, C, E, F, G, H, I, J, K, L, M, N, and O was assessed by Mann-Whitney test and cumulative distribution plots in a by Kolmogorov-Smirnov test. *p < 0.05, ** p < 0.01.

**Figure S7:**
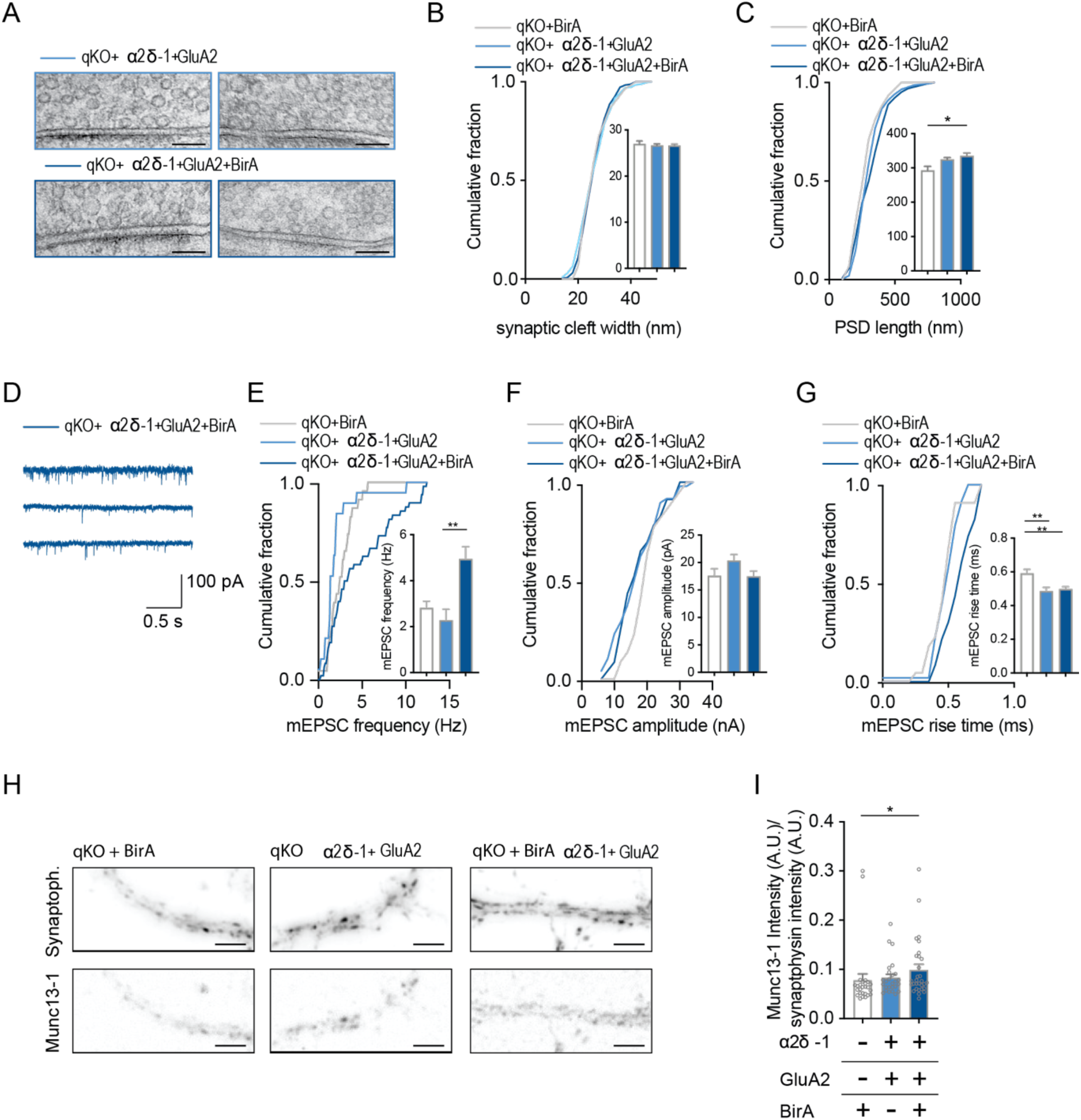
The artificial alignment of α 2δ 1 and GluA2 partially rescues mEPSC frequency and Munc13-1 protein levels. **(A)** Representative electron micrographs of cryofixed primary hippocampal neurons from qKO mice expressing α2δ1-AP and GluA2-mSA or α2δ1-AP, GluA2-mSA, and BirA. Scale bars 100 nm (**B)** Cumulative probability plot and bar graph of the synaptic cleft width. (**C)** Cumulative probability plot and bar graph of the PSD length. (**D)** Example traces of miniature postsynaptic currents (mEPSCs).(**E)** Cumulative probability plot, and bar graph of mEPSC frequency. (**F)** Cumulative probability plot, and bar graph of mEPSC amplitude. (**G)** Cumulative probability plot, and bar graph of mEPSC rise time. (**H)** Example images of Munc13-1 and synaptophysin immunostainings in neuronal cultures obtained from qKO mice. Scale bars 5 µm. (**I)** Munc13-1 immunofluorescence intensities normalized to synaptophysin intensities. Data are means ± SEM. Statistical significance for B, C, E, F, G, and I was assessed by Kruskal-Wallis test. *p < 0.05, **p < 0.01.

**Table S1.**
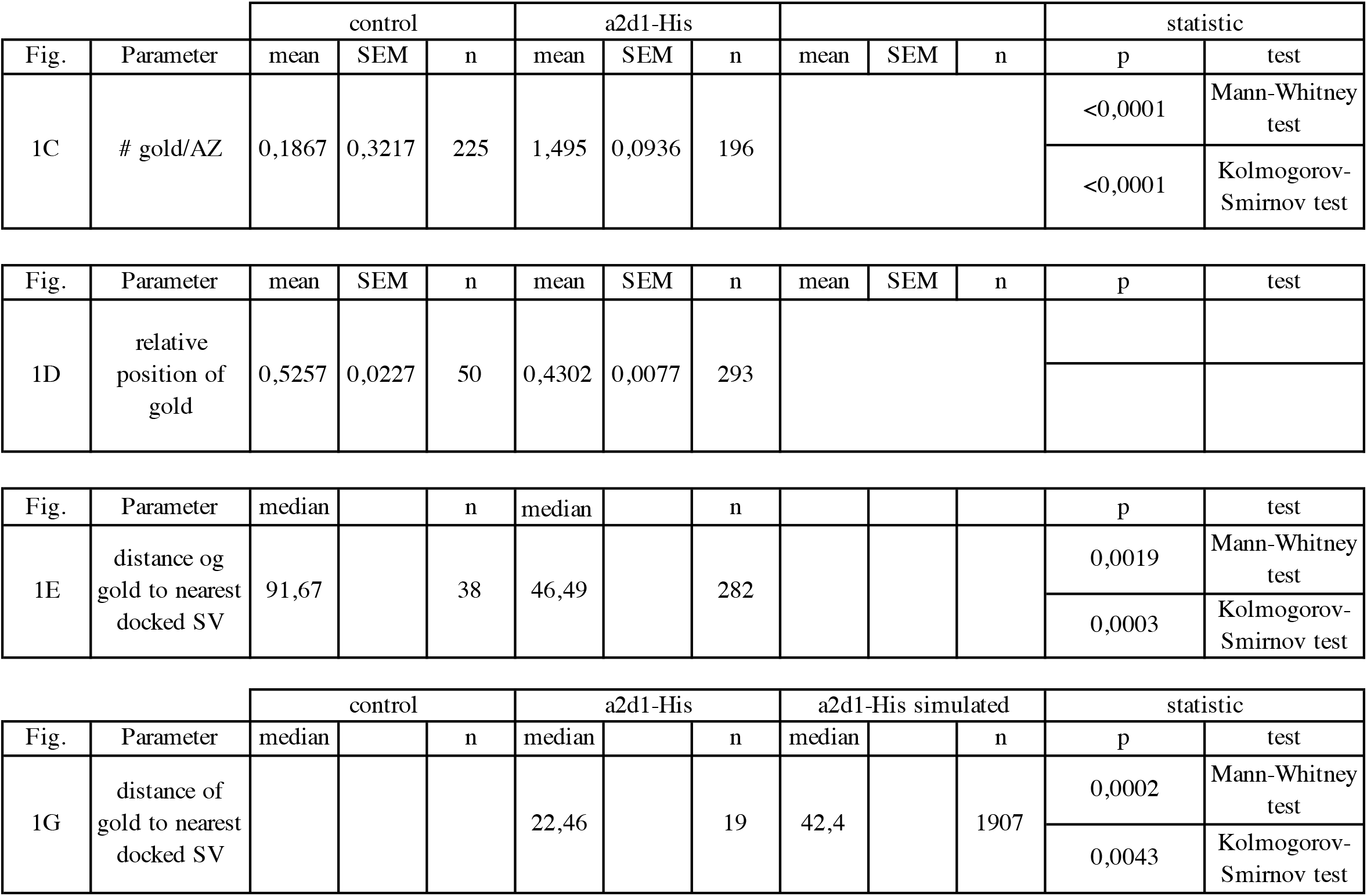

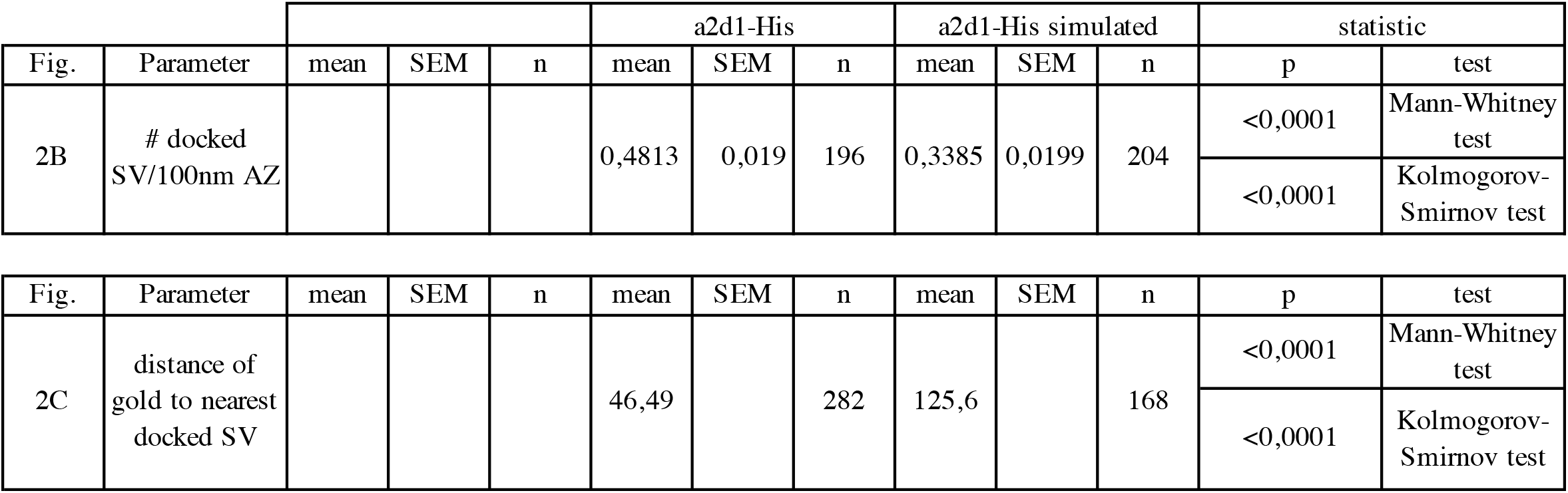

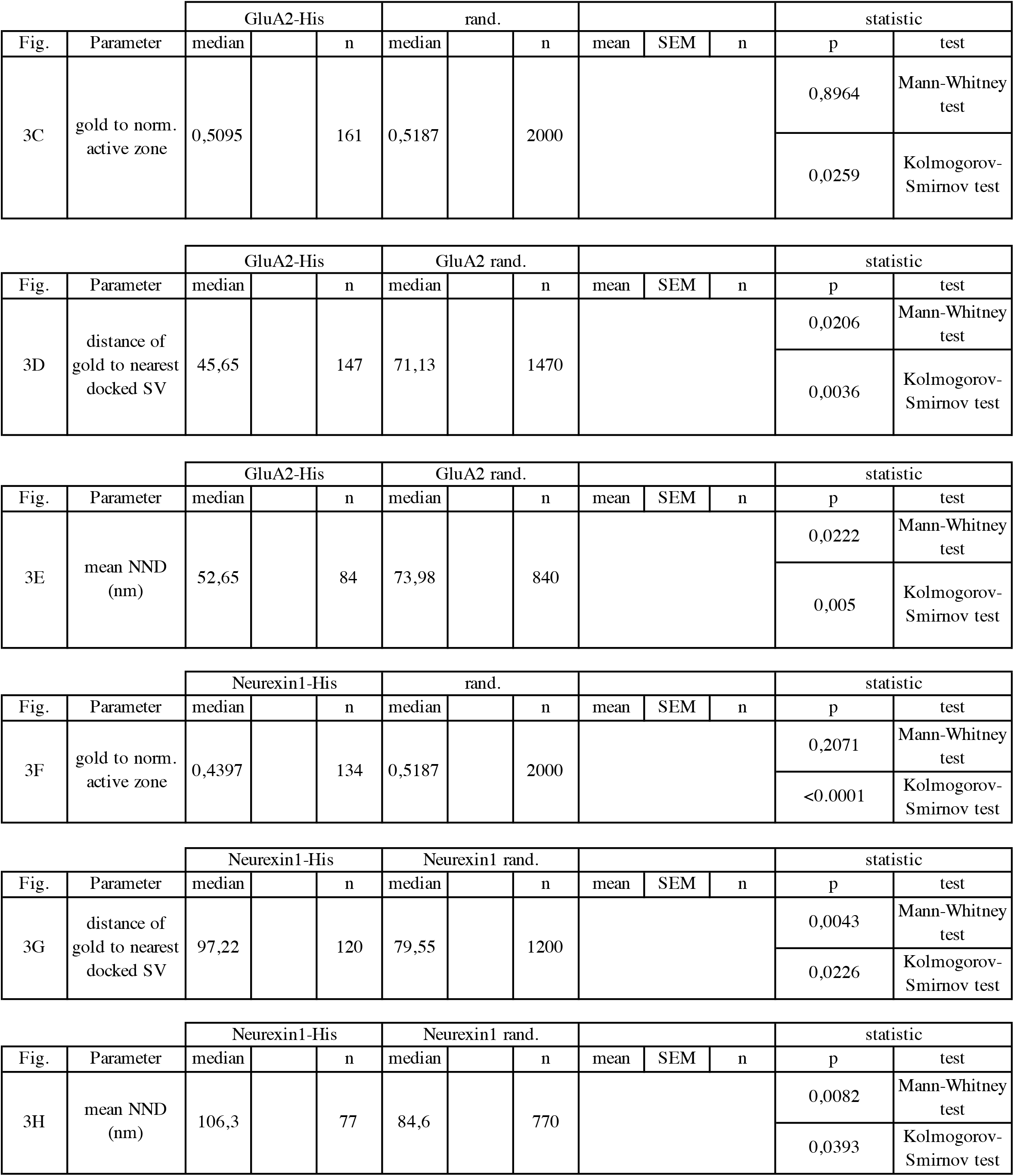

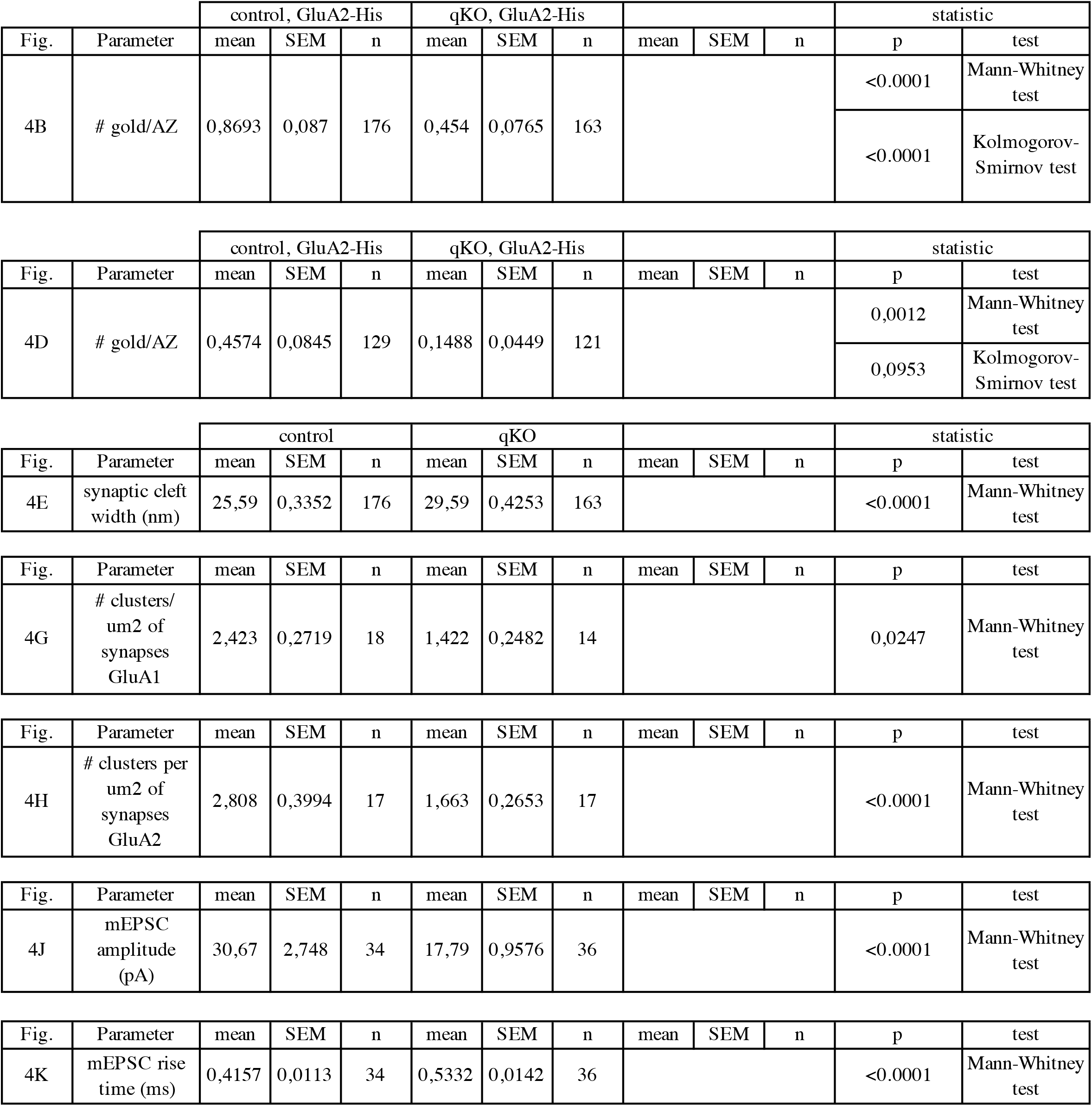

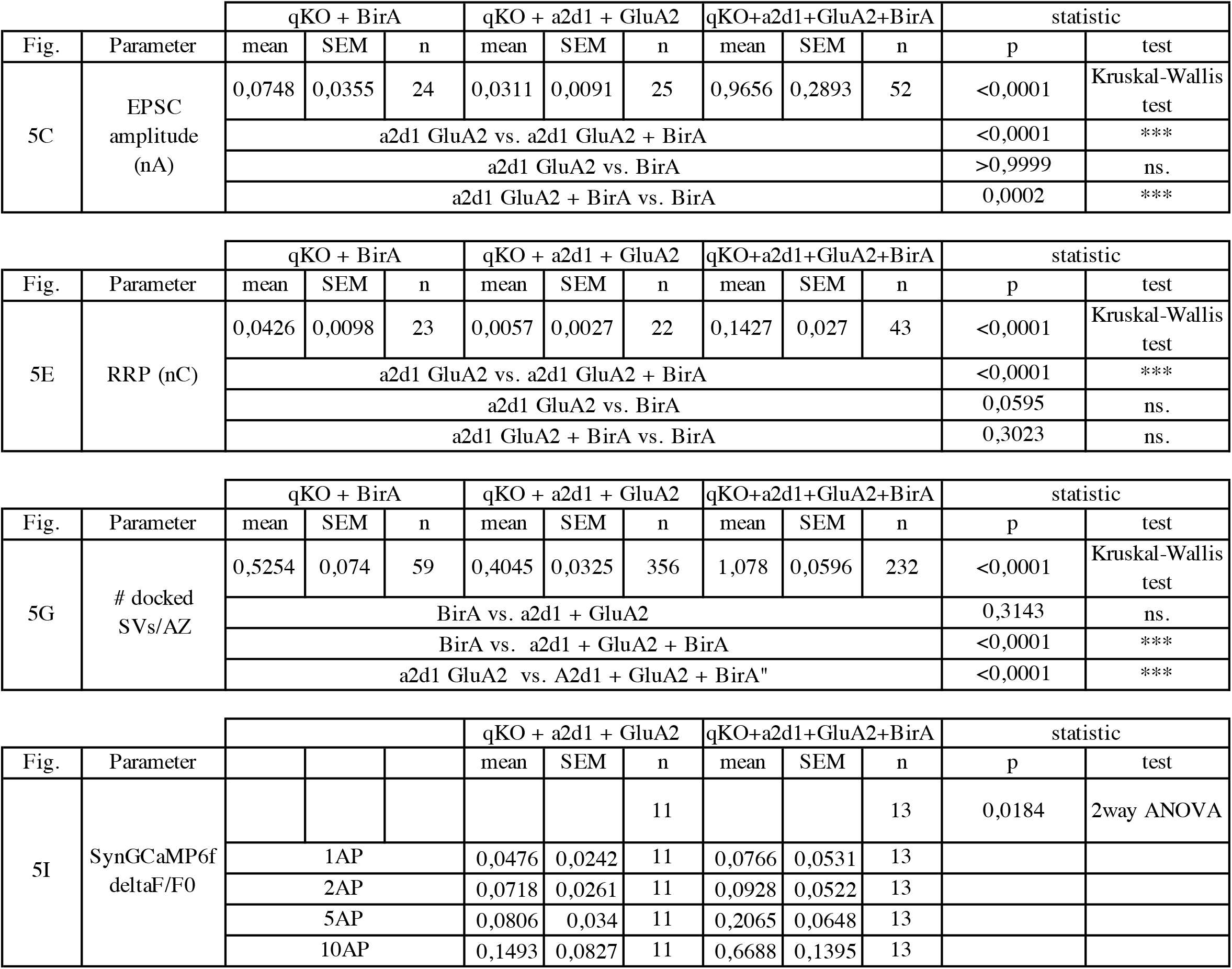

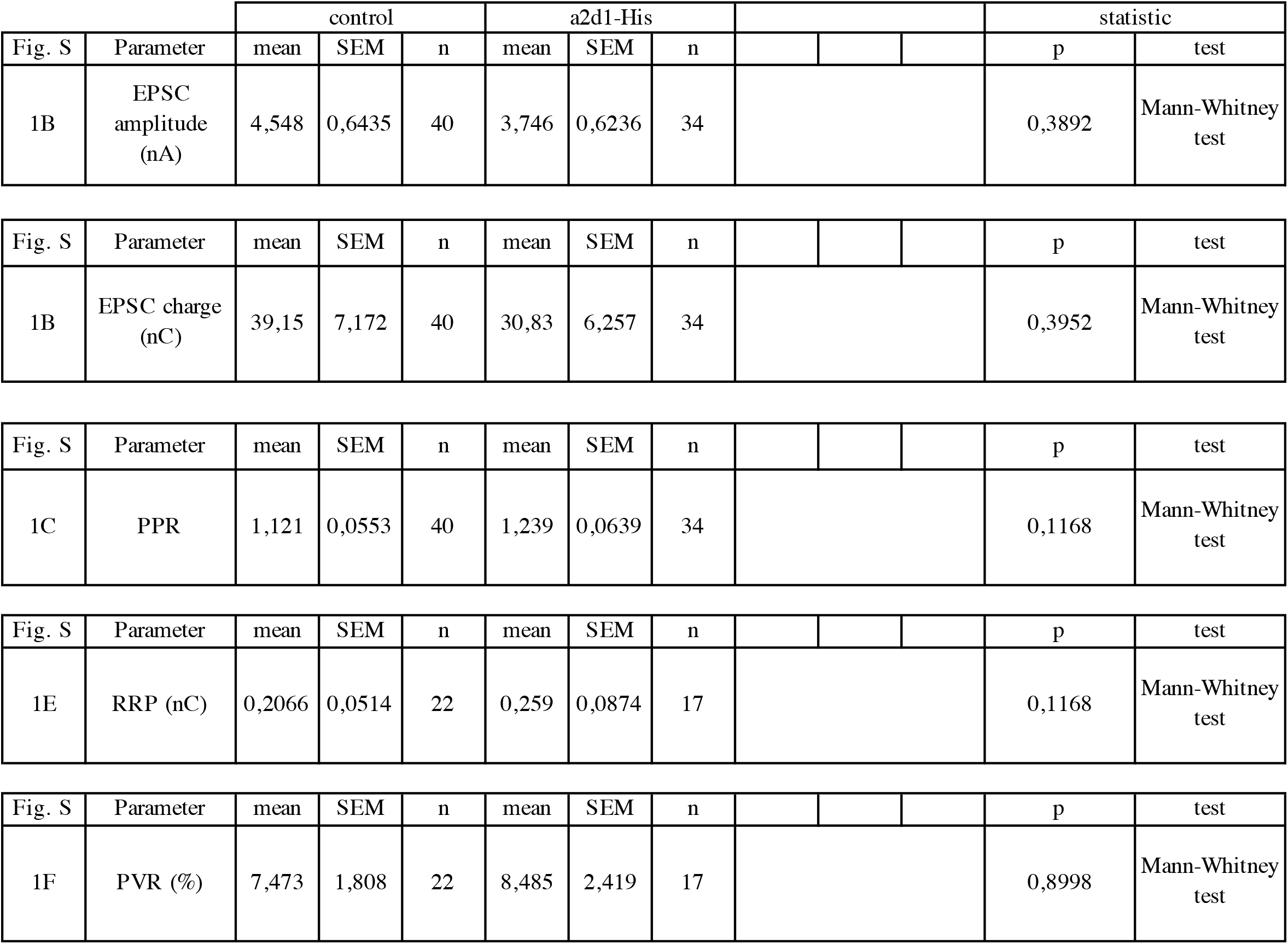

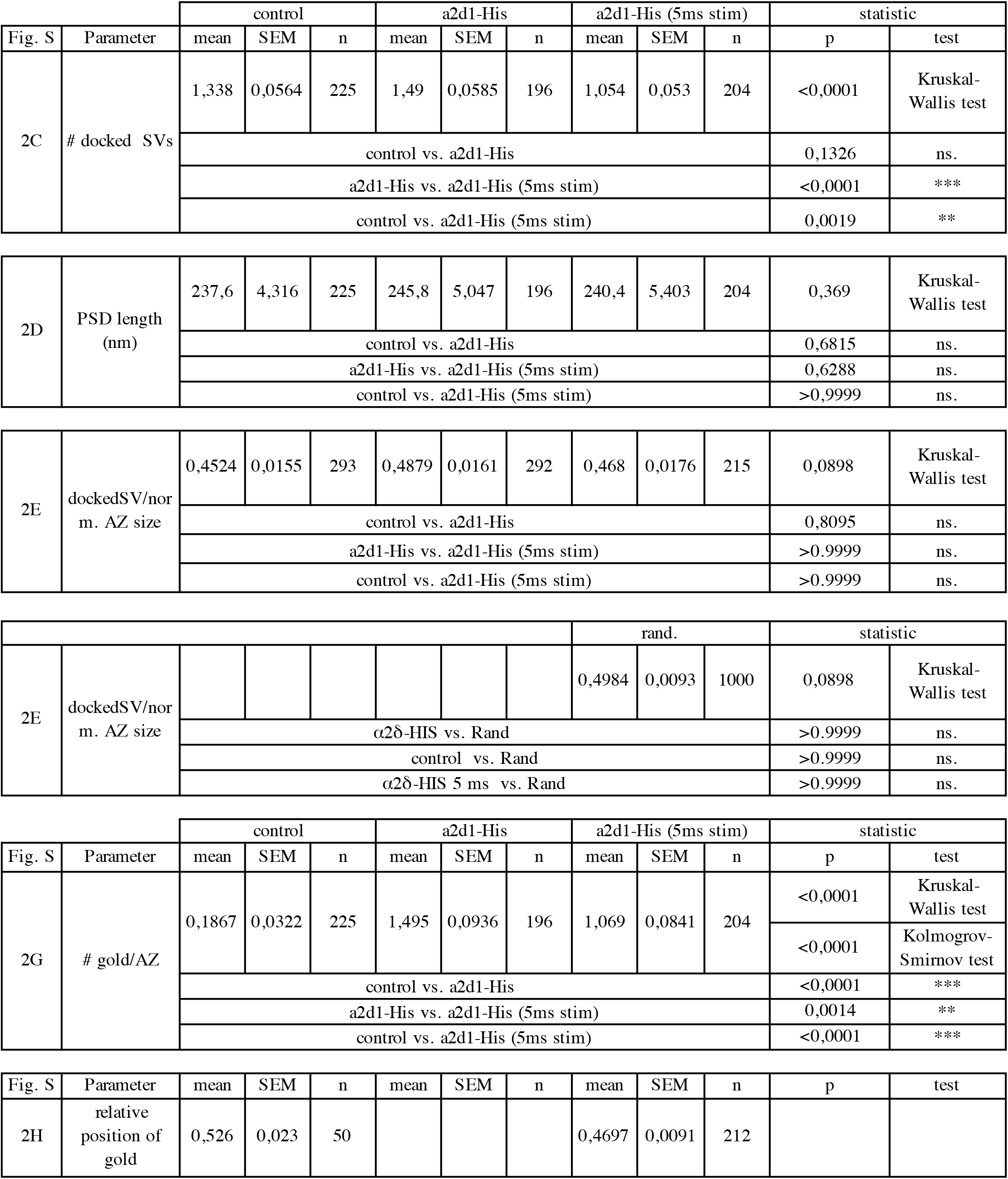

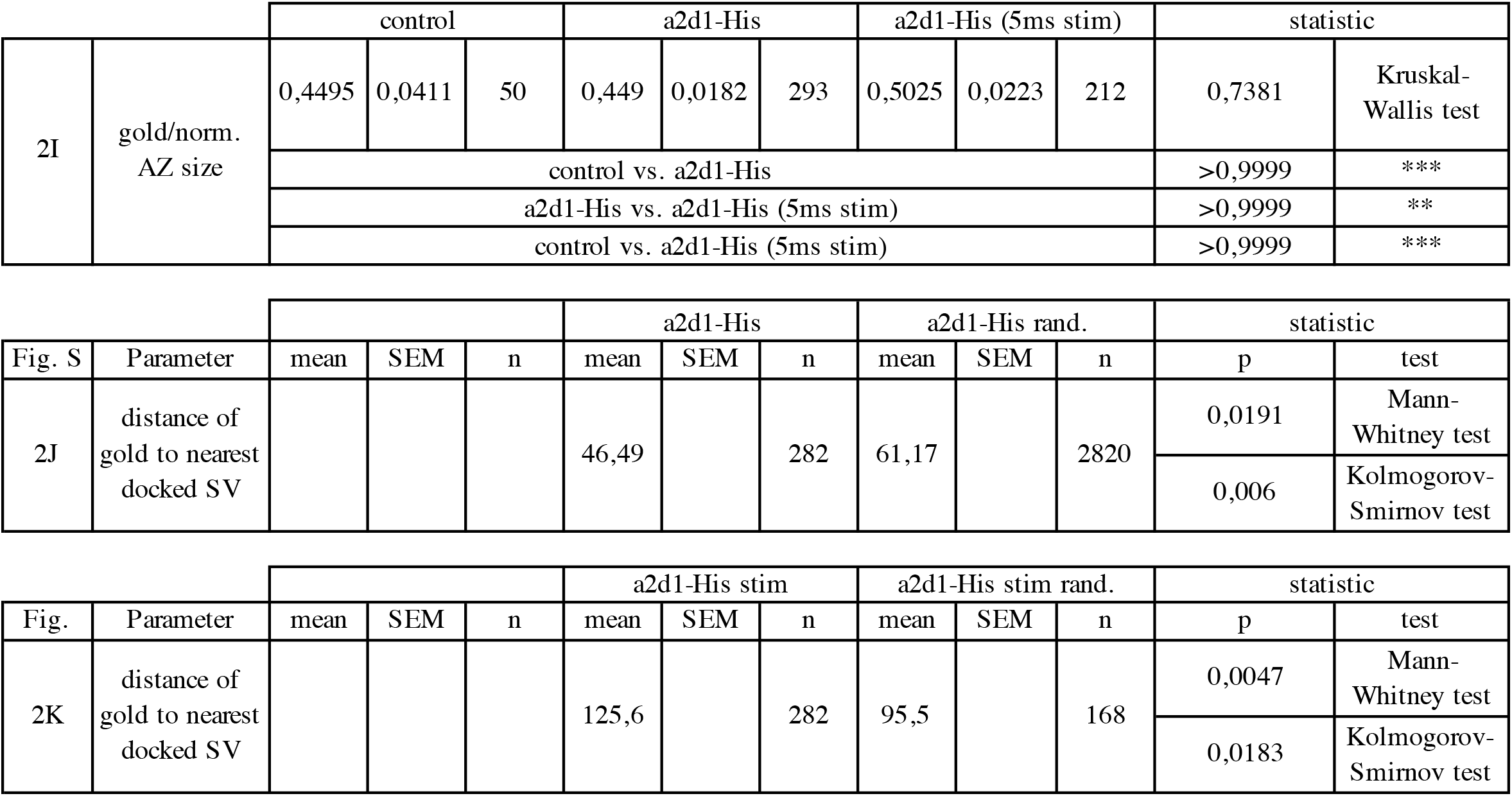

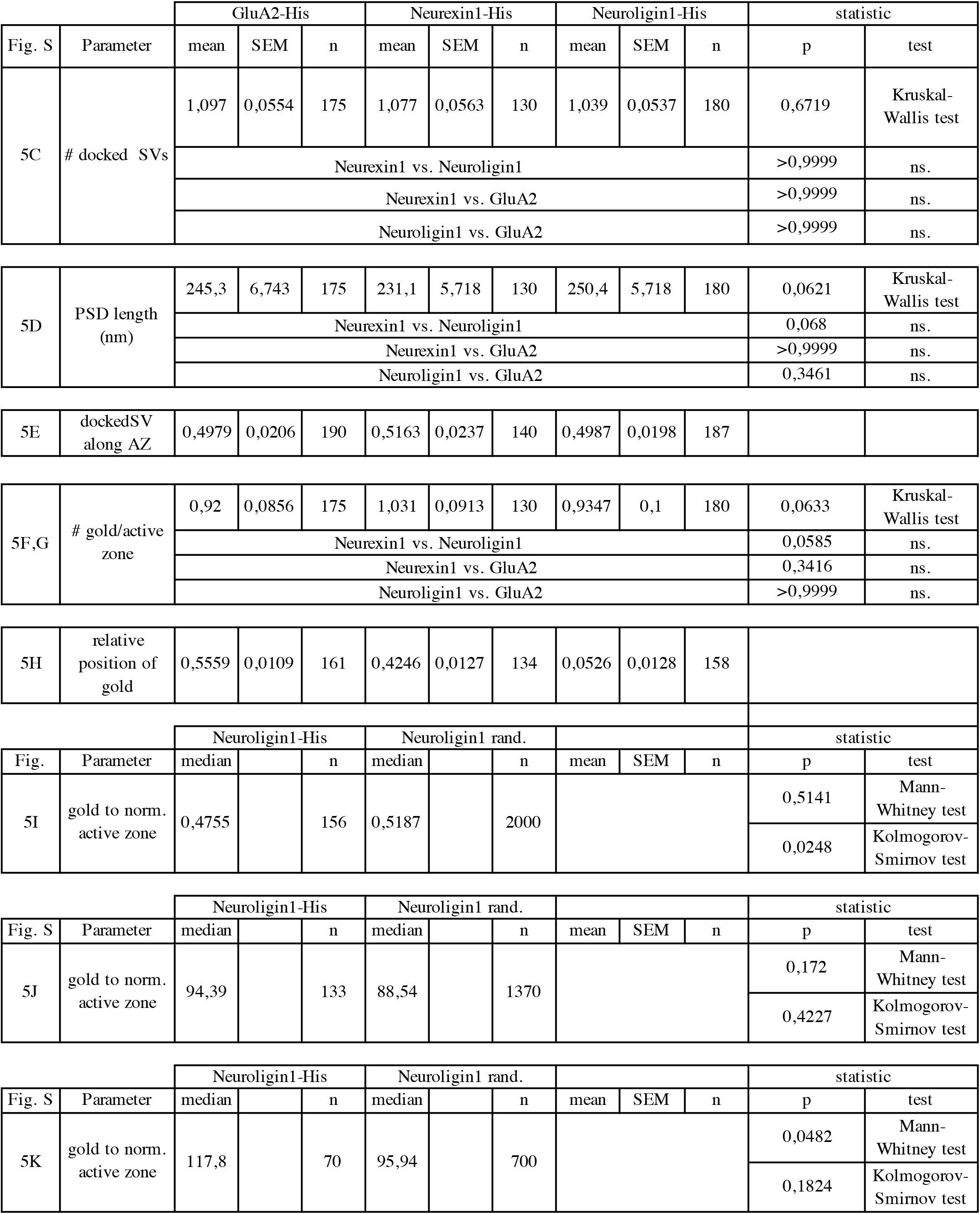

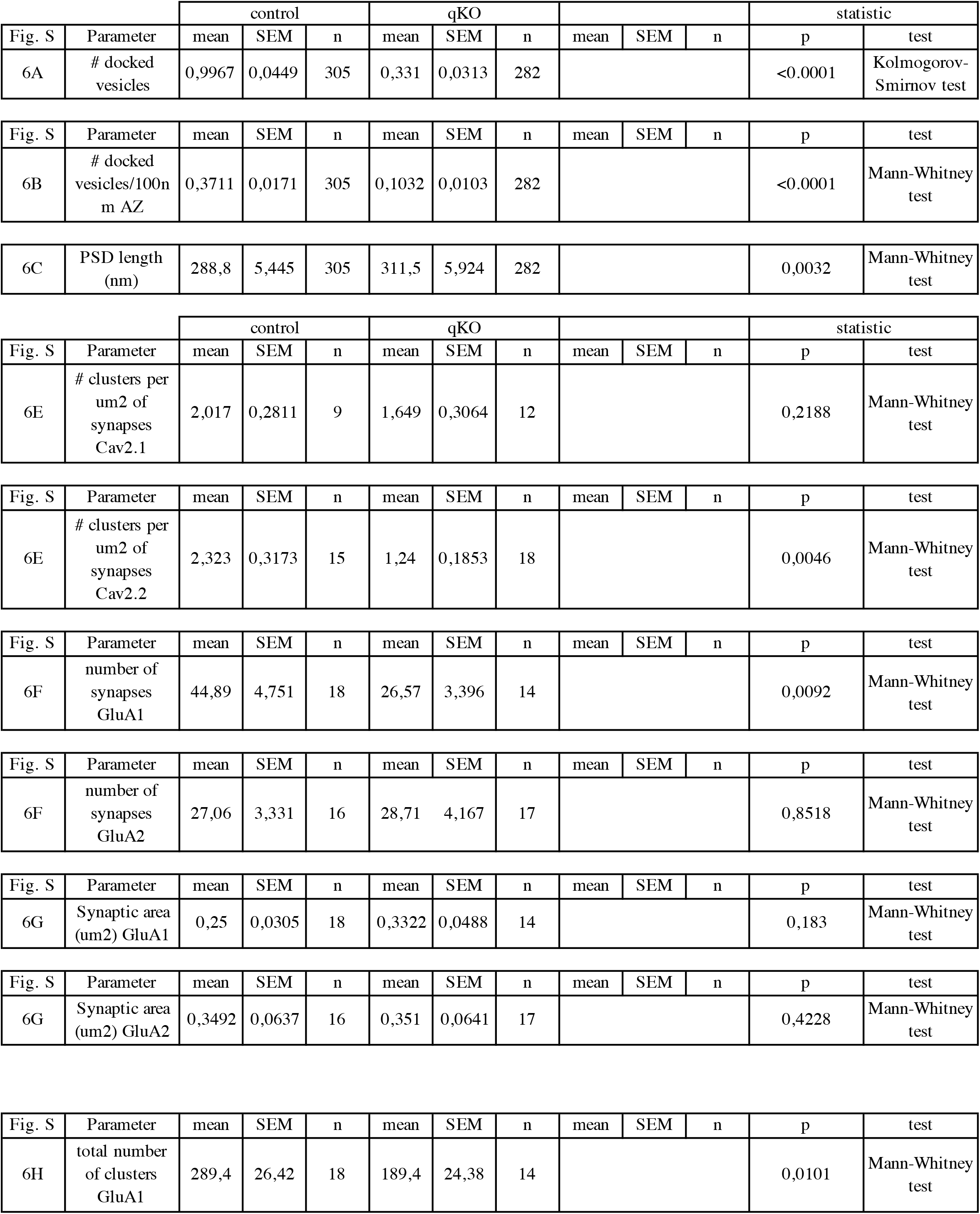
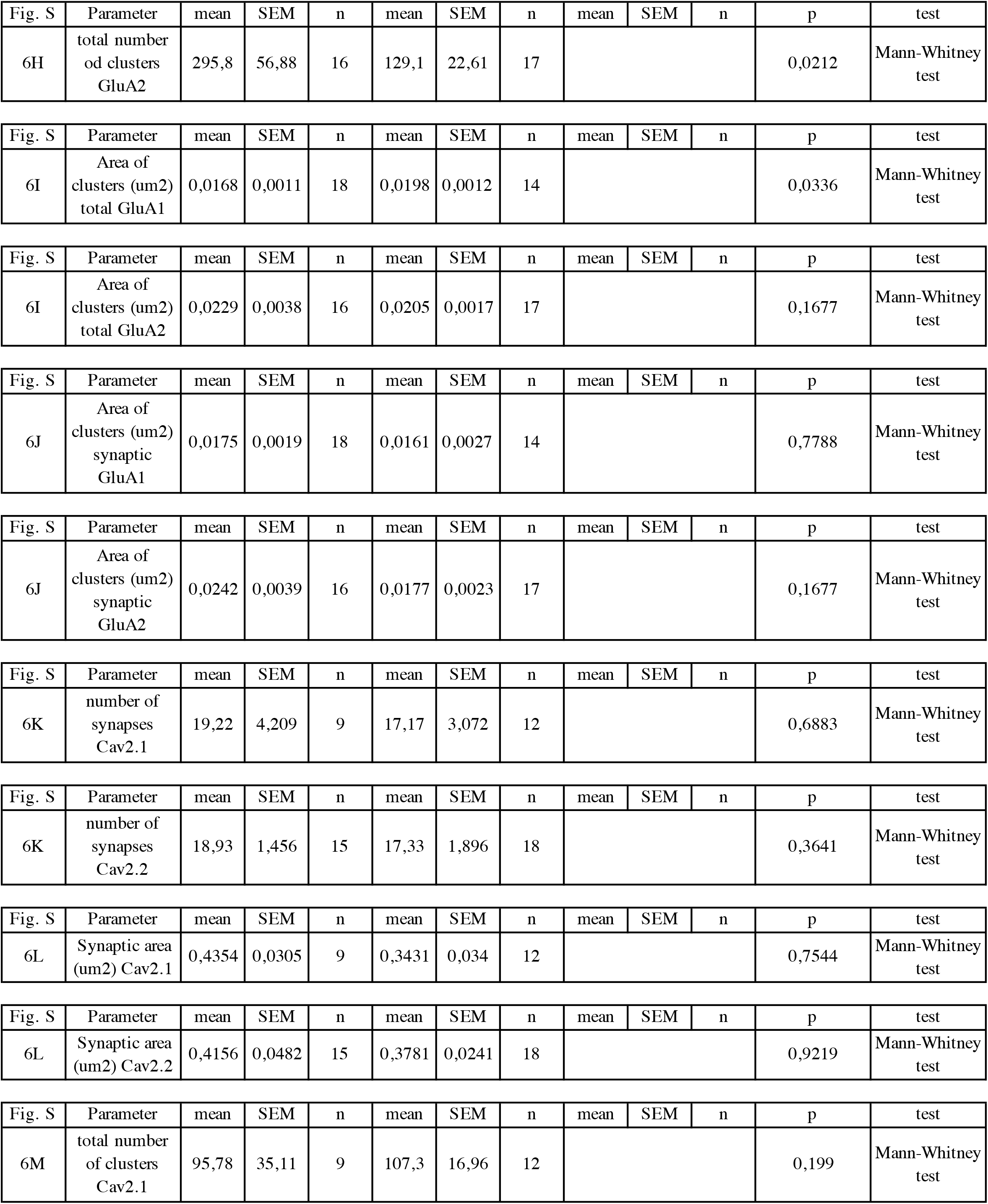
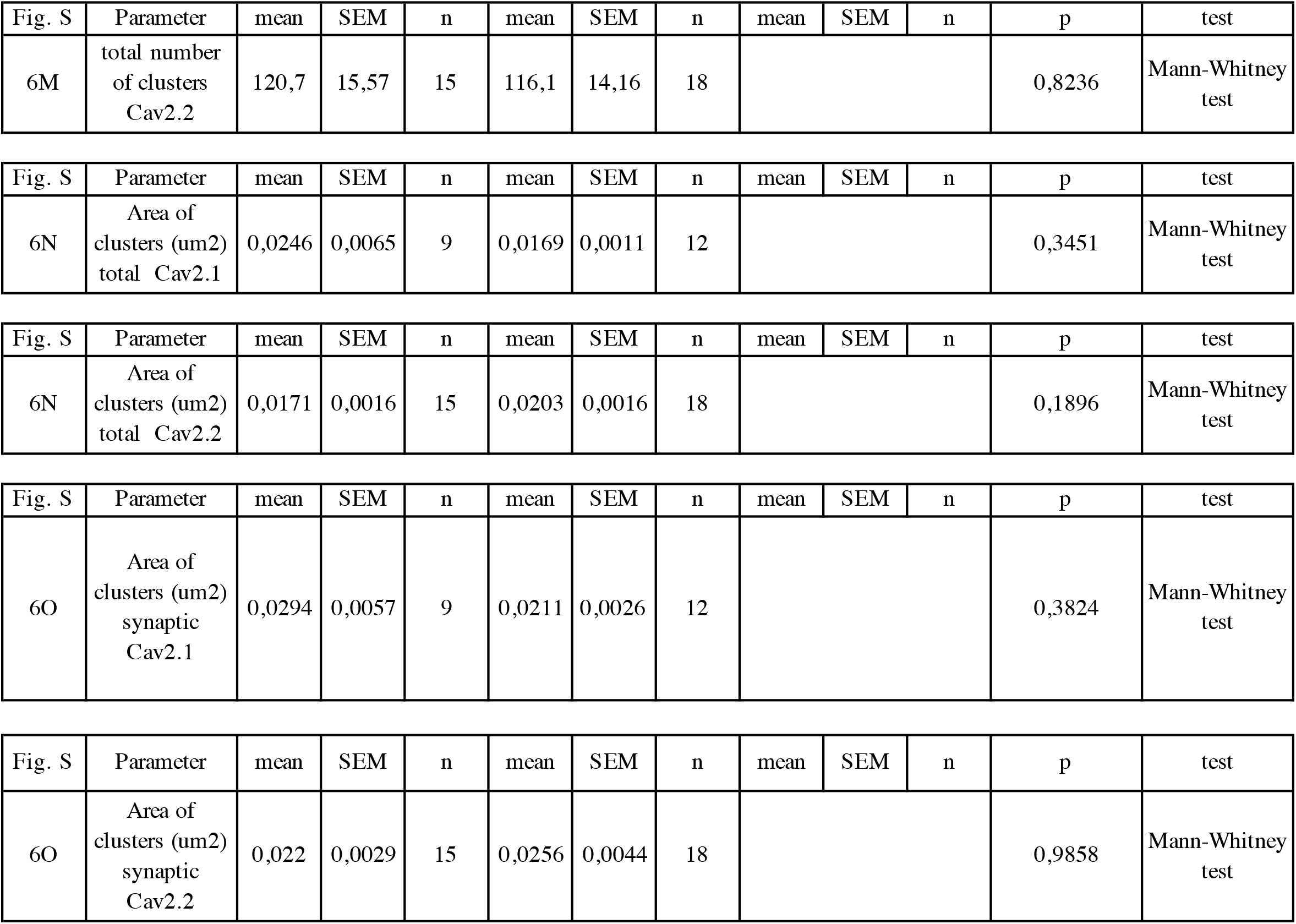

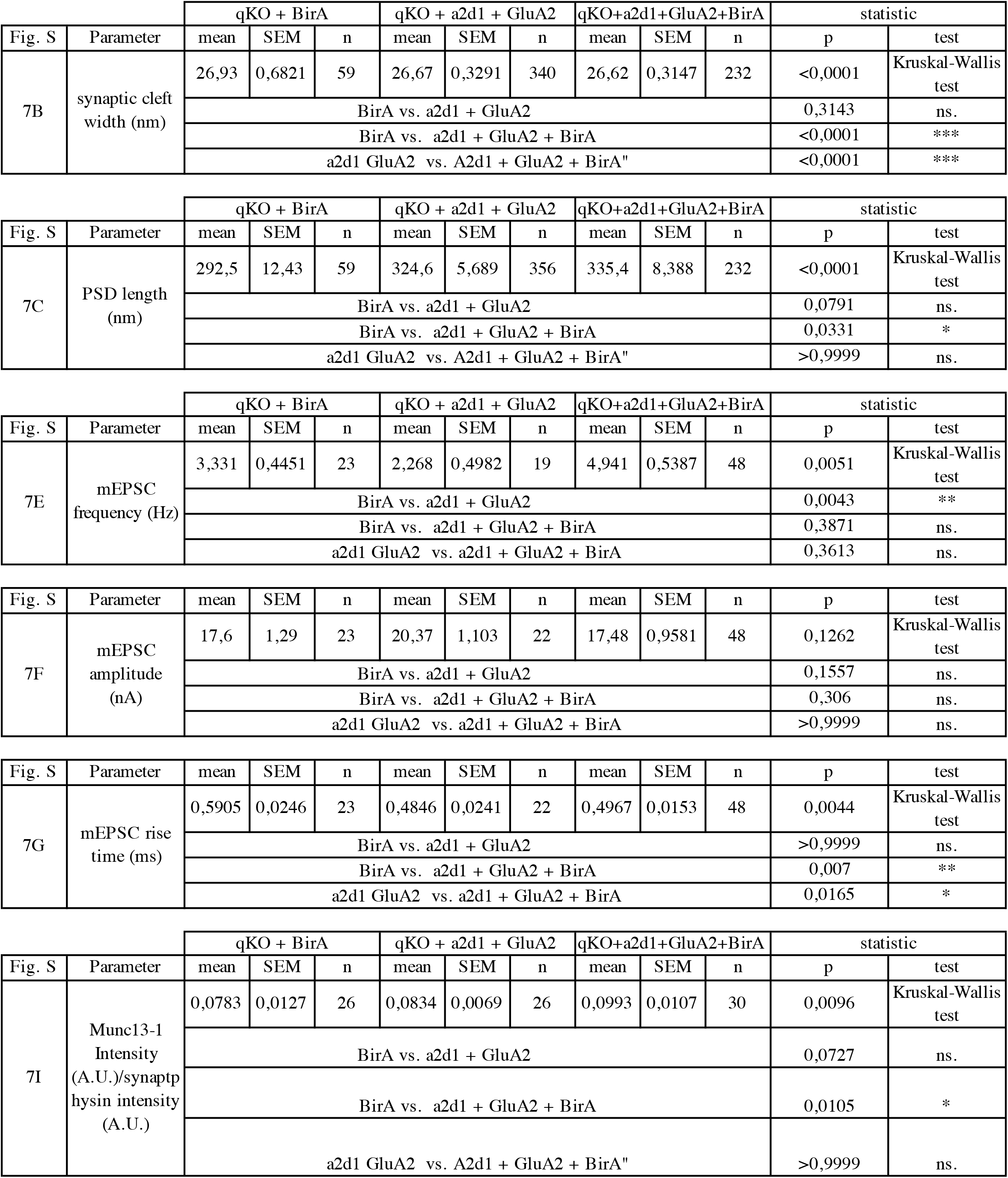

