## Supplementary material for "Functional architecture of the synaptic transducers at a central glutamatergic synapse": Statistics

**Table S1.**

### Figure 1

| | | control | | | a2d1-His | | | | | | statistic | |
|---|---|---|---|---|---|---|---|---|---|---|---|---|
| Fig. | Parameter | mean | SEM | n | mean | SEM | n | mean | SEM | n | p | test |
| 1C | # gold/AZ | 0,1867 | 0,3217 | 225 | 1,495 | 0,0936 | 196 | | | | <0,0001 | Mann-Whitney test |
| | | | | | | | | | | | <0,0001 | Kolmogorov-Smirnov test |

| Fig. | Parameter | mean | SEM | n | mean | SEM | n | mean | SEM | n | p | test |
|---|---|---|---|---|---|---|---|---|---|---|---|---|
| 1D | relative position of gold | 0,5257 | 0,0227 | 50 | 0,4302 | 0,0077 | 293 | | | | | |

| Fig. | Parameter | median | | n | median | | n | | | | p | test |
|---|---|---|---|---|---|---|---|---|---|---|---|---|
| 1E | distance og gold to nearest docked SV | 91,67 | | 38 | 46,49 | | 282 | | | | 0,0019 | Mann-Whitney test |
| | | | | | | | | | | | 0,0003 | Kolmogorov-Smirnov test |

| | | control | | | a2d1-His | | | a2d1-His simulated | | | statistic | |
|---|---|---|---|---|---|---|---|---|---|---|---|---|
| Fig. | Parameter | median | | n | median | | n | median | | n | p | test |
| 1G | distance of gold to nearest docked SV | | | | 22,46 | | 19 | 42,4 | | 1907 | 0,0002 | Mann-Whitney test |
| | | | | | | | | | | | 0,0043 | Kolmogorov-Smirnov test |

**Table S1.**

### Figure 2

| | | | | | a2d1-His | | | a2d1-His simulated | | | statistic | |
|---|---|---|---|---|---|---|---|---|---|---|---|---|
| Fig. | Parameter | mean | SEM | n | mean | SEM | n | mean | SEM | n | p | test |
| 2B | # docked SV/100nm AZ | | | | 0,4813 | 0,019 | 196 | 0,3385 | 0,0199 | 204 | <0,0001 | Mann-Whitney test |
| | | | | | | | | | | | <0,0001 | Kolmogorov-Smirnov test |

| Fig. | Parameter | mean | SEM | n | mean | SEM | n | mean | SEM | n | p | test |
|---|---|---|---|---|---|---|---|---|---|---|---|---|
| 2C | distance of gold to nearest docked SV | | | | 46,49 | | 282 | 125,6 | | 168 | <0,0001 | Mann-Whitney test |
| | | | | | | | | | | | <0,0001 | Kolmogorov-Smirnov test |

### Figure 3

| | | GluA2-His | | | rand. | | | | | | statistic | |
|---|---|---|---|---|---|---|---|---|---|---|---|---|
| Fig. | Parameter | median | | n | median | | n | mean | SEM | n | p | test |
| 3C | gold to norm. active zone | 0,5095 | | 161 | 0,5187 | | 2000 | | | | 0,8964 | Mann-Whitney test |
| | | | | | | | | | | | 0,0259 | Kolmogorov-Smirnov test |

| | | GluA2-His | | | GluA2 rand. | | | | | | statistic | |
|---|---|---|---|---|---|---|---|---|---|---|---|---|
| Fig. | Parameter | median | | n | median | | n | mean | SEM | n | p | test |
| 3D | distance of gold to nearest docked SV | 45,65 | | 147 | 71,13 | | 1470 | | | | 0,0206 | Mann-Whitney test |
| | | | | | | | | | | | 0,0036 | Kolmogorov-Smirnov test |

| | | GluA2-His | | | GluA2 rand. | | | | | | statistic | |
|---|---|---|---|---|---|---|---|---|---|---|---|---|
| Fig. | Parameter | median | | n | median | | n | mean | SEM | n | p | test |
| 3E | mean NND (nm) | 52,65 | | 84 | 73,98 | | 840 | | | | 0,0222 | Mann-Whitney test |
| | | | | | | | | | | | 0,005 | Kolmogorov-Smirnov test |

| | | Neurexin1-His | | | rand. | | | | | | statistic | |
|---|---|---|---|---|---|---|---|---|---|---|---|---|
| Fig. | Parameter | median | | n | median | | n | mean | SEM | n | p | test |
| 3F | gold to norm. active zone | 0,4397 | | 134 | 0,5187 | | 2000 | | | | 0,2071 | Mann-Whitney test |
| | | | | | | | | | | | <0.0001 | Kolmogorov-Smirnov test |

| | | Neurexin1-His | | | Neurexin1 rand. | | | | | | statistic | |
|---|---|---|---|---|---|---|---|---|---|---|---|---|
| Fig. | Parameter | median | | n | median | | n | mean | SEM | n | p | test |
| 3G | distance of gold to nearest docked SV | 97,22 | | 120 | 79,55 | | 1200 | | | | 0,0043 | Mann-Whitney test |
| | | | | | | | | | | | 0,0226 | Kolmogorov-Smirnov test |

| | | Neurexin1-His | | | Neurexin1 rand. | | | | | | statistic | |
|---|---|---|---|---|---|---|---|---|---|---|---|---|
| Fig. | Parameter | median | | n | median | | n | mean | SEM | n | p | test |
| 3H | mean NND (nm) | 106,3 | | 77 | 84,6 | | 770 | | | | 0,0082 | Mann-Whitney test |
| | | | | | | | | | | | 0,0393 | Kolmogorov-Smirnov test |

### Figure 4

| | | control, GluA2-His | | | qKO, GluA2-His | | | | | | statistic | |
|---|---|---|---|---|---|---|---|---|---|---|---|---|
| Fig. | Parameter | mean | SEM | n | mean | SEM | n | mean | SEM | n | p | test |
| 4B | # gold/AZ | 0,8693 | 0,087 | 176 | 0,454 | 0,0765 | 163 | | | | <0.0001 | Mann-Whitney test |
| | | | | | | | | | | | <0.0001 | Kolmogorov-Smirnov test |

| | | control, GluA2-His | | | qKO, GluA2-His | | | | | | statistic | |
|---|---|---|---|---|---|---|---|---|---|---|---|---|
| Fig. | Parameter | mean | SEM | n | mean | SEM | n | mean | SEM | n | p | test |
| 4D | # gold/AZ | 0,4574 | 0,0845 | 129 | 0,1488 | 0,0449 | 121 | | | | 0,0012 | Mann-Whitney test |
| | | | | | | | | | | | 0,0953 | Kolmogorov-Smirnov test |

| | | control | | | qKO | | | | | | statistic | |
|---|---|---|---|---|---|---|---|---|---|---|---|---|
| Fig. | Parameter | mean | SEM | n | mean | SEM | n | mean | SEM | n | p | test |
| 4E | synaptic cleft width (nm) | 25,59 | 0,3352 | 176 | 29,59 | 0,4253 | 163 | | | | <0.0001 | Mann-Whitney test |

| Fig. | Parameter | mean | SEM | n | mean | SEM | n | mean | SEM | n | p | test |
|---|---|---|---|---|---|---|---|---|---|---|---|---|
| 4G | # clusters/ um2 of synapses GluA1 | 2,423 | 0,2719 | 18 | 1,422 | 0,2482 | 14 | | | | 0,0247 | Mann-Whitney test |

| Fig. | Parameter | mean | SEM | n | mean | SEM | n | mean | SEM | n | p | test |
|---|---|---|---|---|---|---|---|---|---|---|---|---|
| 4H | # clusters per um2 of synapses GluA2 | 2,808 | 0,3994 | 17 | 1,663 | 0,2653 | 17 | | | | <0.0001 | Mann-Whitney test |

| Fig. | Parameter | mean | SEM | n | mean | SEM | n | mean | SEM | n | p | test |
|---|---|---|---|---|---|---|---|---|---|---|---|---|
| 4J | mEPSC amplitude (pA) | 30,67 | 2,748 | 34 | 17,79 | 0,9576 | 36 | | | | <0.0001 | Mann-Whitney test |

| Fig. | Parameter | mean | SEM | n | mean | SEM | n | mean | SEM | n | p | test |
|---|---|---|---|---|---|---|---|---|---|---|---|---|
| 4K | mEPSC rise time (ms) | 0,4157 | 0,0113 | 34 | 0,5332 | 0,0142 | 36 | | | | <0.0001 | Mann-Whitney test |

### Figure 5

| | | qKO + BirA | | | qKO + a2d1 + GluA2 | | | qKO+a2d1+GluA2+BirA | | | statistic | |
|---|---|---|---|---|---|---|---|---|---|---|---|---|
| Fig. | Parameter | mean | SEM | n | mean | SEM | n | mean | SEM | n | p | test |
| 5C | EPSC amplitude (nA) | 0,0748 | 0,0355 | 24 | 0,0311 | 0,0091 | 25 | 0,9656 | 0,2893 | 52 | <0,0001 | Kruskal-Wallis test |
| | | a2d1 GluA2 vs. a2d1 GluA2 + BirA | | | | | | | | | <0,0001 | *** |
| | | a2d1 GluA2 vs. BirA | | | | | | | | | >0,9999 | ns. |
| | | a2d1 GluA2 + BirA vs. BirA | | | | | | | | | 0,0002 | *** |

| | | qKO + BirA | | | qKO + a2d1 + GluA2 | | | qKO+a2d1+GluA2+BirA | | | statistic | |
|---|---|---|---|---|---|---|---|---|---|---|---|---|
| Fig. | Parameter | mean | SEM | n | mean | SEM | n | mean | SEM | n | p | test |
| 5E | RRP (nC) | 0,0426 | 0,0098 | 23 | 0,0057 | 0,0027 | 22 | 0,1427 | 0,027 | 43 | <0,0001 | Kruskal-Wallis test |
| | | a2d1 GluA2 vs. a2d1 GluA2 + BirA | | | | | | | | | <0,0001 | *** |
| | | a2d1 GluA2 vs. BirA | | | | | | | | | 0,0595 | ns. |
| | | a2d1 GluA2 + BirA vs. BirA | | | | | | | | | 0,3023 | ns. |

| | | qKO + BirA | | | qKO + a2d1 + GluA2 | | | qKO+a2d1+GluA2+BirA | | | statistic | |
|---|---|---|---|---|---|---|---|---|---|---|---|---|
| Fig. | Parameter | mean | SEM | n | mean | SEM | n | mean | SEM | n | p | test |
| 5G | # docked SVs/AZ | 0,5254 | 0,074 | 59 | 0,4045 | 0,0325 | 356 | 1,078 | 0,0596 | 232 | <0,0001 | Kruskal-Wallis test |
| | | BirA vs. a2d1 + GluA2 | | | | | | | | | 0,3143 | ns. |
| | | BirA vs. a2d1 + GluA2 + BirA | | | | | | | | | <0,0001 | *** |
| | | a2d1 GluA2 vs. A2d1 + GluA2 + BirA" | | | | | | | | | <0,0001 | *** |

| | | | | | qKO + a2d1 + GluA2 | | | qKO+a2d1+GluA2+BirA | | | statistic | |
|---|---|---|---|---|---|---|---|---|---|---|---|---|
| Fig. | Parameter | | | | mean | SEM | n | mean | SEM | n | p | test |
| 5I | SynGCaMP6f deltaF/F0 | | | | | | 11 | | | 13 | 0,0184 | 2way ANOVA |
| | | 1AP | | | 0,0476 | 0,0242 | 11 | 0,0766 | 0,0531 | 13 | | |
| | | 2AP | | | 0,0718 | 0,0261 | 11 | 0,0928 | 0,0522 | 13 | | |
| | | 5AP | | | 0,0806 | 0,034 | 11 | 0,2065 | 0,0648 | 13 | | |
| | | 10AP | | | 0,1493 | 0,0827 | 11 | 0,6688 | 0,1395 | 13 | | |

### Figure S1

| | | control | | | a2d1-His | | | | | | statistic | |
|---|---|---|---|---|---|---|---|---|---|---|---|---|
| Fig. S | Parameter | mean | SEM | n | mean | SEM | n | | | | p | test |
| 1B | EPSC amplitude (nA) | 4,548 | 0,6435 | 40 | 3,746 | 0,6236 | 34 | | | | 0,3892 | Mann-Whitney test |

| Fig. S | Parameter | mean | SEM | n | mean | SEM | n | | | | p | test |
|---|---|---|---|---|---|---|---|---|---|---|---|---|
| 1B | EPSC charge (nC) | 39,15 | 7,172 | 40 | 30,83 | 6,257 | 34 | | | | 0,3952 | Mann-Whitney test |

| Fig. S | Parameter | mean | SEM | n | mean | SEM | n | | | | p | test |
|---|---|---|---|---|---|---|---|---|---|---|---|---|
| 1C | PPR | 1,121 | 0,0553 | 40 | 1,239 | 0,0639 | 34 | | | | 0,1168 | Mann-Whitney test |

| Fig. S | Parameter | mean | SEM | n | mean | SEM | n | | | | p | test |
|---|---|---|---|---|---|---|---|---|---|---|---|---|
| 1E | RRP (nC) | 0,2066 | 0,0514 | 22 | 0,259 | 0,0874 | 17 | | | | 0,1168 | Mann-Whitney test |

| Fig. S | Parameter | mean | SEM | n | mean | SEM | n | | | | p | test |
|---|---|---|---|---|---|---|---|---|---|---|---|---|
| 1F | PVR (%) | 7,473 | 1,808 | 22 | 8,485 | 2,419 | 17 | | | | 0,8998 | Mann-Whitney test |

### Figure S2

| | | control | | | a2d1-His | | | a2d1-His (5ms stim) | | | statistic | |
|---|---|---|---|---|---|---|---|---|---|---|---|---|
| Fig. S | Parameter | mean | SEM | n | mean | SEM | n | mean | SEM | n | p | test |
| 2C | # docked SVs | 1,338 | 0,0564 | 225 | 1,49 | 0,0585 | 196 | 1,054 | 0,053 | 204 | <0,0001 | Kruskal-Wallis test |
| | | control vs. a2d1-His | | | | | | | | | 0,1326 | ns. |
| | | a2d1-His vs. a2d1-His (5ms stim) | | | | | | | | | <0,0001 | *** |
| | | control vs. a2d1-His (5ms stim) | | | | | | | | | 0,0019 | ** |

| | | | | | | | | | | | | |
|---|---|---|---|---|---|---|---|---|---|---|---|---|
| 2D | PSD length (nm) | 237,6 | 4,316 | 225 | 245,8 | 5,047 | 196 | 240,4 | 5,403 | 204 | 0,369 | Kruskal-Wallis test |
| | | control vs. a2d1-His | | | | | | | | | 0,6815 | ns. |
| | | a2d1-His vs. a2d1-His (5ms stim) | | | | | | | | | 0,6288 | ns. |
| | | control vs. a2d1-His (5ms stim) | | | | | | | | | >0,9999 | ns. |

| | | | | | | | | | | | | |
|---|---|---|---|---|---|---|---|---|---|---|---|---|
| 2E | dockedSV/norm. AZ size | 0,4524 | 0,0155 | 293 | 0,4879 | 0,0161 | 292 | 0,468 | 0,0176 | 215 | 0,0898 | Kruskal-Wallis test |
| | | control vs. a2d1-His | | | | | | | | | 0,8095 | ns. |
| | | a2d1-His vs. a2d1-His (5ms stim) | | | | | | | | | >0.9999 | ns. |
| | | control vs. a2d1-His (5ms stim) | | | | | | | | | >0.9999 | ns. |

| | | | | | | | | rand. | | | statistic | |
|---|---|---|---|---|---|---|---|---|---|---|---|---|
| 2E | dockedSV/norm. AZ size | | | | | | | 0,4984 | 0,0093 | 1000 | 0,0898 | Kruskal-Wallis test |
| | | α2δ-HIS vs. Rand | | | | | | | | | >0.9999 | ns. |
| | | control vs. Rand | | | | | | | | | >0.9999 | ns. |
| | | α2δ-HIS 5 ms vs. Rand | | | | | | | | | >0.9999 | ns. |

| | | control | | | a2d1-His | | | a2d1-His (5ms stim) | | | statistic | |
|---|---|---|---|---|---|---|---|---|---|---|---|---|
| Fig. S | Parameter | mean | SEM | n | mean | SEM | n | mean | SEM | n | p | test |
| 2G | # gold/AZ | 0,1867 | 0,0322 | 225 | 1,495 | 0,0936 | 196 | 1,069 | 0,0841 | 204 | <0,0001 | Kruskal-Wallis test |
| | | | | | | | | | | | <0,0001 | Kolmogrov-Smirnov test |
| | | control vs. a2d1-His | | | | | | | | | <0,0001 | *** |
| | | a2d1-His vs. a2d1-His (5ms stim) | | | | | | | | | 0,0014 | ** |
| | | control vs. a2d1-His (5ms stim) | | | | | | | | | <0,0001 | *** |

| Fig. S | Parameter | mean | SEM | n | mean | SEM | n | mean | SEM | n | p | test |
|---|---|---|---|---|---|---|---|---|---|---|---|---|
| 2H | relative position of gold | 0,526 | 0,023 | 50 | | | | 0,4697 | 0,0091 | 212 | | |

### Figure S2

| | | control | | | a2d1-His | | | a2d1-His (5ms stim) | | | statistic | |
|---|---|---|---|---|---|---|---|---|---|---|---|---|
| 2I | gold/norm. AZ size | 0,4495 | 0,0411 | 50 | 0,449 | 0,0182 | 293 | 0,5025 | 0,0223 | 212 | 0,7381 | Kruskal-Wallis test |
| | | control vs. a2d1-His | | | | | | | | | >0,9999 | *** |
| | | a2d1-His vs. a2d1-His (5ms stim) | | | | | | | | | >0,9999 | ** |
| | | control vs. a2d1-His (5ms stim) | | | | | | | | | >0,9999 | *** |

| | | | | | a2d1-His | | | a2d1-His rand. | | | statistic | |
|---|---|---|---|---|---|---|---|---|---|---|---|---|
| Fig. S | Parameter | mean | SEM | n | mean | SEM | n | mean | SEM | n | p | test |
| 2J | distance of gold to nearest docked SV | | | | 46,49 | | 282 | 61,17 | | 2820 | 0,0191 | Mann-Whitney test |
| | | | | | | | | | | | 0,006 | Kolmogorov-Smirnov test |

| | | | | | a2d1-His stim | | | a2d1-His stim rand. | | | statistic | |
|---|---|---|---|---|---|---|---|---|---|---|---|---|
| Fig. | Parameter | mean | SEM | n | mean | SEM | n | mean | SEM | n | p | test |
| 2K | distance of gold to nearest docked SV | | | | 125,6 | | 282 | 95,5 | | 168 | 0,0047 | Mann-Whitney test |
| | | | | | | | | | | | 0,0183 | Kolmogorov-Smirnov test |

### Figure S5

| Fig. S | Parameter | GluA2-His | | | Neurexin1-His | | | Neuroligin1-His | | | statistic | |
|---|---|---|---|---|---|---|---|---|---|---|---|---|
| | | mean | SEM | n | mean | SEM | n | mean | SEM | n | p | test |
| 5C | # docked SVs | 1,097 | 0,0554 | 175 | 1,077 | 0,0563 | 130 | 1,039 | 0,0537 | 180 | 0,6719 | Kruskal-Wallis test |
| | | Neurexin1 vs. Neuroligin1 | | | | | | | | | >0,9999 | ns. |
| | | Neurexin1 vs. GluA2 | | | | | | | | | >0,9999 | ns. |
| | | Neuroligin1 vs. GluA2 | | | | | | | | | >0,9999 | ns. |
| 5D | PSD length (nm) | 245,3 | 6,743 | 175 | 231,1 | 5,718 | 130 | 250,4 | 5,718 | 180 | 0,0621 | Kruskal-Wallis test |
| | | Neurexin1 vs. Neuroligin1 | | | | | | | | | 0,068 | ns. |
| | | Neurexin1 vs. GluA2 | | | | | | | | | >0,9999 | ns. |
| | | Neuroligin1 vs. GluA2 | | | | | | | | | 0,3461 | ns. |
| 5E | dockedSV along AZ | 0,4979 | 0,0206 | 190 | 0,5163 | 0,0237 | 140 | 0,4987 | 0,0198 | 187 | | |
| 5F,G | # gold/active zone | 0,92 | 0,0856 | 175 | 1,031 | 0,0913 | 130 | 0,9347 | 0,1 | 180 | 0,0633 | Kruskal-Wallis test |
| | | Neurexin1 vs. Neuroligin1 | | | | | | | | | 0,0585 | ns. |
| | | Neurexin1 vs. GluA2 | | | | | | | | | 0,3416 | ns. |
| | | Neuroligin1 vs. GluA2 | | | | | | | | | >0,9999 | ns. |
| 5H | relative position of gold | 0,5559 | 0,0109 | 161 | 0,4246 | 0,0127 | 134 | 0,0526 | 0,0128 | 158 | | |

| Fig. | Parameter | Neuroligin1-His | | | Neuroligin1 rand. | | | | | | statistic | |
|---|---|---|---|---|---|---|---|---|---|---|---|---|
| | | median | | n | median | | n | mean | SEM | n | p | test |
| 5I | gold to norm. active zone | 0,4755 | | 156 | 0,5187 | | 2000 | | | | 0,5141 | Mann-Whitney test |
| | | | | | | | | | | | 0,0248 | Kolmogorov-Smirnov test |

| Fig. S | Parameter | Neuroligin1-His | | | Neuroligin1 rand. | | | | | | statistic | |
|---|---|---|---|---|---|---|---|---|---|---|---|---|
| | | median | | n | median | | n | mean | SEM | n | p | test |
| 5J | gold to norm. active zone | 94,39 | | 133 | 88,54 | | 1370 | | | | 0,172 | Mann-Whitney test |
| | | | | | | | | | | | 0,4227 | Kolmogorov-Smirnov test |

| Fig. S | Parameter | Neuroligin1-His | | | Neuroligin1 rand. | | | | | | statistic | |
|---|---|---|---|---|---|---|---|---|---|---|---|---|
| | | median | | n | median | | n | mean | SEM | n | p | test |
| 5K | gold to norm. active zone | 117,8 | | 70 | 95,94 | | 700 | | | | 0,0482 | Mann-Whitney test |
| | | | | | | | | | | | 0,1824 | Kolmogorov-Smirnov test |

### Figure S6

| | | control | | | qKO | | | | | | statistic | |
|---|---|---|---|---|---|---|---|---|---|---|---|---|
| Fig. S | Parameter | mean | SEM | n | mean | SEM | n | mean | SEM | n | p | test |
| 6A | # docked vesicles | 0,9967 | 0,0449 | 305 | 0,331 | 0,0313 | 282 | | | | <0.0001 | Kolmogorov-Smirnov test |

| Fig. S | Parameter | mean | SEM | n | mean | SEM | n | mean | SEM | n | p | test |
|---|---|---|---|---|---|---|---|---|---|---|---|---|
| 6B | # docked vesicles/100nm AZ | 0,3711 | 0,0171 | 305 | 0,1032 | 0,0103 | 282 | | | | <0.0001 | Mann-Whitney test |
| 6C | PSD length (nm) | 288,8 | 5,445 | 305 | 311,5 | 5,924 | 282 | | | | 0,0032 | Mann-Whitney test |

| | | control | | | qKO | | | | | | statistic | |
|---|---|---|---|---|---|---|---|---|---|---|---|---|
| Fig. S | Parameter | mean | SEM | n | mean | SEM | n | mean | SEM | n | p | test |
| 6E | # clusters per um2 of synapses Cav2.1 | 2,017 | 0,2811 | 9 | 1,649 | 0,3064 | 12 | | | | 0,2188 | Mann-Whitney test |

| Fig. S | Parameter | mean | SEM | n | mean | SEM | n | mean | SEM | n | p | test |
|---|---|---|---|---|---|---|---|---|---|---|---|---|
| 6E | # clusters per um2 of synapses Cav2.2 | 2,323 | 0,3173 | 15 | 1,24 | 0,1853 | 18 | | | | 0,0046 | Mann-Whitney test |

| Fig. S | Parameter | mean | SEM | n | mean | SEM | n | mean | SEM | n | p | test |
|---|---|---|---|---|---|---|---|---|---|---|---|---|
| 6F | number of synapses GluA1 | 44,89 | 4,751 | 18 | 26,57 | 3,396 | 14 | | | | 0,0092 | Mann-Whitney test |

| Fig. S | Parameter | mean | SEM | n | mean | SEM | n | mean | SEM | n | p | test |
|---|---|---|---|---|---|---|---|---|---|---|---|---|
| 6F | number of synapses GluA2 | 27,06 | 3,331 | 16 | 28,71 | 4,167 | 17 | | | | 0,8518 | Mann-Whitney test |

| Fig. S | Parameter | mean | SEM | n | mean | SEM | n | mean | SEM | n | p | test |
|---|---|---|---|---|---|---|---|---|---|---|---|---|
| 6G | Synaptic area (um2) GluA1 | 0,25 | 0,0305 | 18 | 0,3322 | 0,0488 | 14 | | | | 0,183 | Mann-Whitney test |

| Fig. S | Parameter | mean | SEM | n | mean | SEM | n | mean | SEM | n | p | test |
|---|---|---|---|---|---|---|---|---|---|---|---|---|
| 6G | Synaptic area (um2) GluA2 | 0,3492 | 0,0637 | 16 | 0,351 | 0,0641 | 17 | | | | 0,4228 | Mann-Whitney test |

| Fig. S | Parameter | mean | SEM | n | mean | SEM | n | mean | SEM | n | p | test |
|---|---|---|---|---|---|---|---|---|---|---|---|---|
| 6H | total number of clusters GluA1 | 289,4 | 26,42 | 18 | 189,4 | 24,38 | 14 | | | | 0,0101 | Mann-Whitney test |

### Figure S6

| Fig. S | Parameter | mean | SEM | n | mean | SEM | n | mean | SEM | n | p | test |
|---|---|---|---|---|---|---|---|---|---|---|---|---|
| 6H | total number od clusters GluA2 | 295,8 | 56,88 | 16 | 129,1 | 22,61 | 17 | | | | 0,0212 | Mann-Whitney test |

| Fig. S | Parameter | mean | SEM | n | mean | SEM | n | mean | SEM | n | p | test |
|---|---|---|---|---|---|---|---|---|---|---|---|---|
| 6I | Area of clusters (um2) total GluA1 | 0,0168 | 0,0011 | 18 | 0,0198 | 0,0012 | 14 | | | | 0,0336 | Mann-Whitney test |

| Fig. S | Parameter | mean | SEM | n | mean | SEM | n | mean | SEM | n | p | test |
|---|---|---|---|---|---|---|---|---|---|---|---|---|
| 6I | Area of clusters (um2) total GluA2 | 0,0229 | 0,0038 | 16 | 0,0205 | 0,0017 | 17 | | | | 0,1677 | Mann-Whitney test |

| Fig. S | Parameter | mean | SEM | n | mean | SEM | n | mean | SEM | n | p | test |
|---|---|---|---|---|---|---|---|---|---|---|---|---|
| 6J | Area of clusters (um2) synaptic GluA1 | 0,0175 | 0,0019 | 18 | 0,0161 | 0,0027 | 14 | | | | 0,7788 | Mann-Whitney test |

| Fig. S | Parameter | mean | SEM | n | mean | SEM | n | mean | SEM | n | p | test |
|---|---|---|---|---|---|---|---|---|---|---|---|---|
| 6J | Area of clusters (um2) synaptic GluA2 | 0,0242 | 0,0039 | 16 | 0,0177 | 0,0023 | 17 | | | | 0,1677 | Mann-Whitney test |

| Fig. S | Parameter | mean | SEM | n | mean | SEM | n | mean | SEM | n | p | test |
|---|---|---|---|---|---|---|---|---|---|---|---|---|
| 6K | number of synapses Cav2.1 | 19,22 | 4,209 | 9 | 17,17 | 3,072 | 12 | | | | 0,6883 | Mann-Whitney test |

| Fig. S | Parameter | mean | SEM | n | mean | SEM | n | mean | SEM | n | p | test |
|---|---|---|---|---|---|---|---|---|---|---|---|---|
| 6K | number of synapses Cav2.2 | 18,93 | 1,456 | 15 | 17,33 | 1,896 | 18 | | | | 0,3641 | Mann-Whitney test |

| Fig. S | Parameter | mean | SEM | n | mean | SEM | n | mean | SEM | n | p | test |
|---|---|---|---|---|---|---|---|---|---|---|---|---|
| 6L | Synaptic area (um2) Cav2.1 | 0,4354 | 0,0305 | 9 | 0,3431 | 0,034 | 12 | | | | 0,7544 | Mann-Whitney test |

| Fig. S | Parameter | mean | SEM | n | mean | SEM | n | mean | SEM | n | p | test |
|---|---|---|---|---|---|---|---|---|---|---|---|---|
| 6L | Synaptic area (um2) Cav2.2 | 0,4156 | 0,0482 | 15 | 0,3781 | 0,0241 | 18 | | | | 0,9219 | Mann-Whitney test |

| Fig. S | Parameter | mean | SEM | n | mean | SEM | n | mean | SEM | n | p | test |
|---|---|---|---|---|---|---|---|---|---|---|---|---|
| 6M | total number of clusters Cav2.1 | 95,78 | 35,11 | 9 | 107,3 | 16,96 | 12 | | | | 0,199 | Mann-Whitney test |

### Figure S6

| Fig. S | Parameter | mean | SEM | n | mean | SEM | n | mean | SEM | n | p | test |
|---|---|---|---|---|---|---|---|---|---|---|---|---|
| 6M | total number of clusters Cav2.2 | 120,7 | 15,57 | 15 | 116,1 | 14,16 | 18 | | | | 0,8236 | Mann-Whitney test |

| Fig. S | Parameter | mean | SEM | n | mean | SEM | n | mean | SEM | n | p | test |
|---|---|---|---|---|---|---|---|---|---|---|---|---|
| 6N | Area of clusters (um2) total Cav2.1 | 0,0246 | 0,0065 | 9 | 0,0169 | 0,0011 | 12 | | | | 0,3451 | Mann-Whitney test |

| Fig. S | Parameter | mean | SEM | n | mean | SEM | n | mean | SEM | n | p | test |
|---|---|---|---|---|---|---|---|---|---|---|---|---|
| 6N | Area of clusters (um2) total Cav2.2 | 0,0171 | 0,0016 | 15 | 0,0203 | 0,0016 | 18 | | | | 0,1896 | Mann-Whitney test |

| Fig. S | Parameter | mean | SEM | n | mean | SEM | n | mean | SEM | n | p | test |
|---|---|---|---|---|---|---|---|---|---|---|---|---|
| 6O | Area of clusters (um2) synaptic Cav2.1 | 0,0294 | 0,0057 | 9 | 0,0211 | 0,0026 | 12 | | | | 0,3824 | Mann-Whitney test |

| Fig. S | Parameter | mean | SEM | n | mean | SEM | n | mean | SEM | n | p | test |
|---|---|---|---|---|---|---|---|---|---|---|---|---|
| 6O | Area of clusters (um2) synaptic Cav2.2 | 0,022 | 0,0029 | 15 | 0,0256 | 0,0044 | 18 | | | | 0,9858 | Mann-Whitney test |

### Figure S7

| | | qKO + BirA | | | qKO + a2d1 + GluA2 | | | qKO+a2d1+GluA2+BirA | | | statistic | |
|---|---|---|---|---|---|---|---|---|---|---|---|---|
| Fig. S | Parameter | mean | SEM | n | mean | SEM | n | mean | SEM | n | p | test |
| 7B | synaptic cleft width (nm) | 26,93 | 0,6821 | 59 | 26,67 | 0,3291 | 340 | 26,62 | 0,3147 | 232 | <0,0001 | Kruskal-Wallis test |
| | | BirA vs. a2d1 + GluA2 | | | | | | | | | 0,3143 | ns. |
| | | BirA vs. a2d1 + GluA2 + BirA | | | | | | | | | <0,0001 | *** |
| | | a2d1 GluA2 vs. A2d1 + GluA2 + BirA" | | | | | | | | | <0,0001 | *** |

| | | qKO + BirA | | | qKO + a2d1 + GluA2 | | | qKO+a2d1+GluA2+BirA | | | statistic | |
|---|---|---|---|---|---|---|---|---|---|---|---|---|
| Fig. S | Parameter | mean | SEM | n | mean | SEM | n | mean | SEM | n | p | test |
| 7C | PSD length (nm) | 292,5 | 12,43 | 59 | 324,6 | 5,689 | 356 | 335,4 | 8,388 | 232 | <0,0001 | Kruskal-Wallis test |
| | | BirA vs. a2d1 + GluA2 | | | | | | | | | 0,0791 | ns. |
| | | BirA vs. a2d1 + GluA2 + BirA | | | | | | | | | 0,0331 | * |
| | | a2d1 GluA2 vs. A2d1 + GluA2 + BirA" | | | | | | | | | >0,9999 | ns. |

| | | qKO + BirA | | | qKO + a2d1 + GluA2 | | | qKO+a2d1+GluA2+BirA | | | statistic | |
|---|---|---|---|---|---|---|---|---|---|---|---|---|
| Fig. S | Parameter | mean | SEM | n | mean | SEM | n | mean | SEM | n | p | test |
| 7E | mEPSC frequency (Hz) | 3,331 | 0,4451 | 23 | 2,268 | 0,4982 | 19 | 4,941 | 0,5387 | 48 | 0,0051 | Kruskal-Wallis test |
| | | BirA vs. a2d1 + GluA2 | | | | | | | | | 0,0043 | ** |
| | | BirA vs. a2d1 + GluA2 + BirA | | | | | | | | | 0,3871 | ns. |
| | | a2d1 GluA2 vs. a2d1 + GluA2 + BirA | | | | | | | | | 0,3613 | ns. |

| Fig. S | Parameter | mean | SEM | n | mean | SEM | n | mean | SEM | n | p | test |
|---|---|---|---|---|---|---|---|---|---|---|---|---|
| 7F | mEPSC amplitude (nA) | 17,6 | 1,29 | 23 | 20,37 | 1,103 | 22 | 17,48 | 0,9581 | 48 | 0,1262 | Kruskal-Wallis test |
| | | BirA vs. a2d1 + GluA2 | | | | | | | | | 0,1557 | ns. |
| | | BirA vs. a2d1 + GluA2 + BirA | | | | | | | | | 0,306 | ns. |
| | | a2d1 GluA2 vs. a2d1 + GluA2 + BirA | | | | | | | | | >0,9999 | ns. |

| Fig. S | Parameter | mean | SEM | n | mean | SEM | n | mean | SEM | n | p | test |
|---|---|---|---|---|---|---|---|---|---|---|---|---|
| 7G | mEPSC rise time (ms) | 0,5905 | 0,0246 | 23 | 0,4846 | 0,0241 | 22 | 0,4967 | 0,0153 | 48 | 0,0044 | Kruskal-Wallis test |
| | | BirA vs. a2d1 + GluA2 | | | | | | | | | >0,9999 | ns. |
| | | BirA vs. a2d1 + GluA2 + BirA | | | | | | | | | 0,007 | ** |
| | | a2d1 GluA2 vs. a2d1 + GluA2 + BirA | | | | | | | | | 0,0165 | * |

| | | qKO + BirA | | | qKO + a2d1 + GluA2 | | | qKO+a2d1+GluA2+BirA | | | statistic | |
|---|---|---|---|---|---|---|---|---|---|---|---|---|
| Fig. S | Parameter | mean | SEM | n | mean | SEM | n | mean | SEM | n | p | test |
| 7I | Munc13-1 Intensity (A.U.)/synaptphysin intensity (A.U.) | 0,0783 | 0,0127 | 26 | 0,0834 | 0,0069 | 26 | 0,0993 | 0,0107 | 30 | 0,0096 | Kruskal-Wallis test |
| | | BirA vs. a2d1 + GluA2 | | | | | | | | | 0,0727 | ns. |
| | | BirA vs. a2d1 + GluA2 + BirA | | | | | | | | | 0,0105 | * |
| | | a2d1 GluA2 vs. A2d1 + GluA2 + BirA" | | | | | | | | | >0,9999 | ns. |
